# Accelerated Bayesian inference of population size history from recombining sequence data

**DOI:** 10.1101/2024.03.25.586640

**Authors:** Jonathan Terhorst

## Abstract

I present phlash, a new Bayesian method for inferring population history from whole genome sequence data. phlash is **p**opulation **h**istory **l**earning by **a**veraging **s**ampled **h**istories: it works by drawing random, low-dimensional projections of the coalescent intensity function from the posterior distribution of a psmc-like model, and averaging them together to form an accurate and adaptive size history estimator. On simulated data, phlash tends to be faster and have lower error than several competing methods including smc++, msmc2, and FitCoal. Moreover, it provides a full posterior distribution over population size history, leading to automatic uncertainty quantification of the point estimates, as well to new Bayesian testing procedures for detecting population structure and ancient bottlenecks. On the technical side, the key advance is a novel algorithm for computing the score function (gradient of the log-likelihood) of a coalescent hidden Markov model: when there are *M* hidden states, the algorithm requires. 𝒪(*M* ^2^) time and. 𝒪(1) memory per decoded position, the same cost as evaluating the log-likelihood itself using the naïve forward algorithm. This algorithm is combined with a hand-tuned implementation that fully leverages the power of modern GPU hardware, and the entire method has been released as an easy-to-use Python software package.

## 1 Introduction

Many natural populations experience significant changes in abundance over the course of their existence. Our own species, for example, has increased over a thousand-fold since the advent of agriculture 12,000 years ago, with various human subpopulations having expanded or contracted due to migration, disease, climate change, interbreeding, and other factors (Cavalli-Sforza, 2000; Diamond, 2005). In general, while some change in population size may be due to random chance, measurable growth and decline over evolutionary timescales is frequently the result of interesting biological, cultural, or ecological phenomena. By developing tools to estimate population history from genetic data, a pursuit known as *demographic inference*, we may hope to learn more about the past, and possible future, of our biome.

That said, estimating population history can be lamentably difficult. Signals of this history are only faintly manifested as patterns of allele sharing across sampled individuals. These patterns can be further obscured by natural phenomena such as meiotic recombination or natural selection, or by bioinformatic error. Complex mathematical models are needed to relate the data to a hypothesized size history. Solving these models is computationally expensive, particularly when a large number of samples is analyzed. Moreover, because of a somewhat diffuse relationship between the model and the data, there can be many evolutionary histories that explain a given collection of observations equally well.

Nevertheless, given the potential to enhance our understanding of evolution, significant effort has been invested in developing accurate and user-friendly methods for inferring population size history. An early and well-known example is the pairwise sequentially Markovian coalescent (psmc; Li and Durbin, 2011), which infers historical effective population size using data from a single diploid individual. The basic idea of psmc, outlined in more detail below, is to relate variation in the local time to most recent common ancestor (TMRCA) between a pair of homologous chromosomes to historical fluctuations in population size through a hidden Markov model (HMM). psmc continues to be popular, because it is fast, fairly robust, and does not require phased genotypes, which can be difficult to obtain when working with non-human data (Spence et al., 2018; Mather, Traves, and Ho, 2020) However, it is not without limitations. As originally formulated, the method can only analyze data from a single sample, and it assumes a fairly simplistic evolutionary model in which size history changes at only a small number of pre-determined locations. The latter property leads to obvious visual bias in the resulting estimates, which have a “stair-step” appearance, and has other, less obvious consequences for inference as well (Parag and Pybus, 2019; Ki and Terhorst, 2023).

A number of successor methods have been proposed which remove some of these limitations (Sheehan, Harris, and Yun S Song, 2013; Schiffels and Durbin, 2014; Terhorst, J. A. Kamm, and Yun S Song, 2017; Steinrücken et al., 2019; Schiffels and K. Wang, 2020). Building on the basic psmc model, they are able to analyze larger sample sizes and/or more realistic demographic models involving e.g. population structure and admixture. Several Bayesian variants of psmc have also been proposed (Palacios, Wakeley, and Ramachandran, 2015; Henderson et al., 2021; Ki and Terhorst, 2023), though they are not as widely used, owing perhaps to the inherent computational difficulty of Bayesian inference in this setting. In a related vein, a large related class of methods exists which infer demography using the site frequency spectrum (SFS), a highly compressed summary statistic formed from genotype data (Gutenkunst et al., 2009; Excoffier and Foll, 2011; Excoffier, Dupanloup, et al., 2013; Bhaskar, Y. X. R. Wang, and Y. S. Song, 2015; Jouganous et al., 2017; J. A. Kamm, Terhorst, and Yun S Song, 2017; J. A. Kamm, Terhorst, Durbin, et al., 2020; Excoffier, Marchi, et al., 2021; Hu et al., 2023). These methods are fast, in some cases capable of analyzing tens of thousands of samples, but they completely ignore linkage disequilibrium information, which contains rich information about population history.

In this paper I present phlash, a new Bayesian method for inferring size history from recombining sequence data. phlash aims to combine the advantages of many of the methods mentioned above into single, general purpose inference procedure that is simultaneously fast, accurate, able to analyze many samples (thousands), invariant to phasing, capable of utilizing of both linkage and frequency spectrum information, and able to return a full posterior distribution over the inferred size history function. Aesthetically, phlash estimates have an appealing, non-parametric quality which enables them to adapt to variability in the underlying size history without user intervention or fine-tuning. The key advance that propels these innovations is a novel technique for efficiently differentiating the psmc likelihood function, which enables the method to rapidly navigate to areas of high posterior density. Combined with an extremely efficient, GPU-based software implementation, the end result is a method for performing full Bayesian inference of population size history, at speeds that match (or exceed) many of the optimized methods mentioned above.

## 2 Methods

In this section I formally define the inference problem, the statistical model, and the estimation procedure.

The goal of this paper is to estimate the historical effective population size of a panmictic population using genetic variation data. Mathematically, the estimand is a function *η* [0, ∞) →(0, ∞)such that the effective population size *t* generations ago was *N*_*e*_.(*t*) = [2*η* (*t*)]^−1^. (Except where otherwise noted, the variable *t* and the word “time” always refer to “generations before the present”.) phlash uses a coalescent hidden Markov model to infer *η*, from recombining sequence data, an approach pioneered by Li and Durbin (2011). The basic idea is as follows: given *η*, and recombination and mutation rates *p; θ* > 0, and diploid genotype data **g** ∈ {0; 1}^*L*^ encoding whether two homologous chromosomes are the same or different at each of *L* loci, psmc models **g** as generated according to the latent variable model

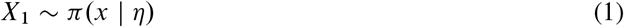

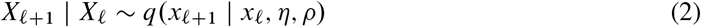

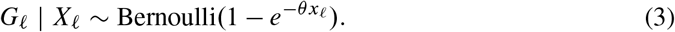

Here, *X*_𝓁_ represents the unobservable time to most recent common ancestor (TMRCA) at genomic position 𝓁, π(*x*| *η*) is its marginal distribution under *η*,, and *q*(*x*_*𝓁*+1_ | *x*_*𝓁*_) is the conditional distribution of TMRCA at a neighboring position given knowledge of the current TMRCA. (The precise form of *q*(· | ·)is not essential here, but see Hobolth and Jensen, 2014, for a detailed investigation.) Note this assumes that the sequence *X*_1_, *X*_2,_ … is Markov. psmc approximates the above generative model by discretizing the time axis, *x*_*𝓁*_ ∈[*t*_*k*,_ *t*_*k*+1_ for some partition

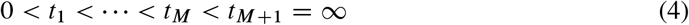

of the real line, and adjusting the transition and emission functions accordingly. A crucial assumption of psmc is that the size history function *η*, also only changes discretely at *t*_*m*_:

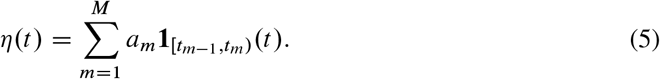

Given these approximations, inference machinery for hidden Markov models can be used to iteratively increase the likelihood of the observed data **g** using the E-M algorithm. The learned parameters are the recombination rate *p*, and the levels *a*_1_, … *a*_*m*_ of *η* in (5). Hence, the choice of discretization (4) thus plays a strong determining role in the overall shape of the estimated function 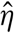 and it is not obvious how these points should be chosen (Terhorst, J. A. Kamm, and Yun S Song, 2017; Ki and Terhorst, 2023). Because of this, in most existing methods, they are selected according to an automated heuristic, with unclear consequences for inference.

Related methods are variations on this approach. Msmc (Schiffels and Durbin, 2014) for example, considers time to first coalescence as the latent state, allowing the analysis of larger sample sizes. Msmc2 (Schiffels and K. Wang, 2020) uses a composite psmc likelihood taken over all pairs of haplotypes. smc++ augments the observed genotype vector **g** with frequency spectrum information, leading to a more complex emission distribution than (3), and optionally uses a spline basis for *η*,. At a high level, though, each of these methods works by discretizing the time axis, as in (4), assuming some parametric representation for *η*, as in (5), and then optimizing to find a point estimate.

### 2.1 Bayesian model

phlash is a simple Bayesian extension of the above model: instead of optimizing to find a single point estimate 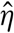 it places a prior on the space of size history functions, and then samples from the posterior distribution

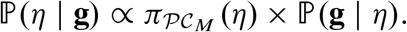

The prior 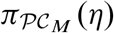 is defined as follows. 𝒫𝒞_***M***_ signifies “piecewise-constant with ***M*** pieces”: let 𝒯_***M***_ be the “ordered tuple space consisting of ordered positive sequences of ***M*** elements,

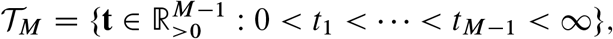

and define 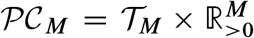. The product space 𝒫𝒞_***M***_ is in bijection with positive, piecewise-constant functions defined on the half-line via the mapping

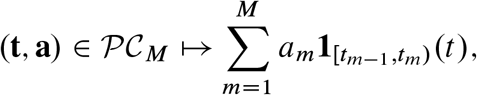

where **1**_*S*_(*x*) denotes the indicator function of the set *S ⊂ ℝ*, and by convention I set *t*_0_ = *0; t*_*M*_ = *∞*. In words, each element (**t, a**) ∈ 𝒫𝒞_***M***_ defines a piecewise-constant function with breaks at *t* = *0, t*_1_, … *t*_*M*-1_, and corresponding levels *a*_1,…,_ *a*_*M*_.

Now I define a prior 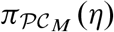 over this space. For all of the examples in this paper, I fixed *M* = 16 and used the following prior:

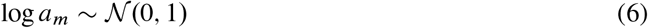

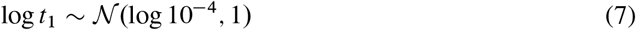

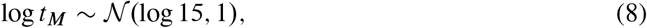

where time is measured in the coalescent scaling, and then set the remaining *t*_*i*_ such that the ratio *t*_*i*+1_*=t*_*i*_ is constant. The choice *M* = 16 is motivated by hardware architecture considerations discussed below.

The key feature of this prior is that the time discretization is random. This means that when a large number of samples from ℙ(*η* | **g**) are averaged together, their individual biases cancel out, resulting in smooth estimates whose shape adapts to the true underlying size history function. Just as important, the user is relieved of the burden of needing to choose the discretization by hand.

### 2.2 Augmenting the model with frequency spectrum information

The psmc model defined above assumes we have a single sample **g**. However, in many modern applications, multiple samples from a population are available, say **G** ∈ {0,1}^*N*x*L*^, where *N* denotes the number of sampled diploids. Additional information from the joint samples can improve estimation accuracy, particularly in the recent past (Schiffels and Durbin, 2014; Terhorst, J. A. Kamm, and Yun S Song, 2017). To enable phlash to analyze larger samples, I define the modified likelihood function

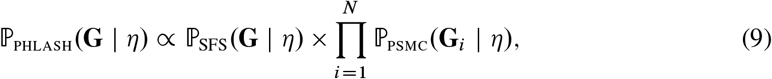

where **G**_*i*_ ∈ {*0; 1*}^*L*^ is the *i* -th row of the genotype matrix **G**. The first term in the product, ℙ_SFS_.(**G** | *η*), is the likelihood of the site frequency spectrum (SFS) obtained from **G** under a “Poisson random field” model (S. A. Sawyer and Hartl, 1992), and can be efficiently computed even for very large samples (Bhaskar, Y. X. R. Wang, and Y. S. Song, 2015). The second term is the product of the psmc likelihood taken over all diploid samples.

Note that the form of (9) is the same as would be obtained if ℙ_SFS_ and ℙ_SFS_.(**G**_*i*_ | *η*) were all pairwise independent, which is not true due to shared ancestry. Thus ℙ_phlash_ is a “composite likelihood”. In general, composite likelihood methods give unbiased point estimates, but overly optimistic confidence intervals if dependence between the component likelihoods is not taken into account (Varin, Reid, and Firth, 2011). The extent to which this affects the dispersion of the posterior distribution returned by phlash is explored in Section 3.4.

### 2.3 Efficient computation of the score function

As with many Bayesian procedures, the principal challenge of the model described above is how to sample from the posterior. State-of-the-art sampling methods like Hamiltonian Monte Carlo (HMC; Neal et al., 2011; Hoffman and Gelman, 2014), stochastic Langevin dynamics (Welling and Teh, 2011), or variational inference (VI; Hoffman, Blei, et al., 2013; Blei, Kucukelbir, and McAuliffe, 2017), all require access to the so-called *score function*, ∇_*η*_log ℙ_phlash_. (*η* |**G)** in the above notation, in order to guide the sampler towards regions of high posterior density. By Bayes’ rule, the score function equals ∇_*η*_log ℙ_phlash_ .(**G**| *η*), so in view of (9) and the fact that ℙ_SFS_. (**G**| *η*), is essentially just a matrix-vector multiply (Polanski and Kimmel, 2003), the main obstacle to sampling is being able to efficiently calculate ∇_*η*_log ℙ_psmc_ (**g** | *η*).

#### 2.3.1 Novel algorithm

The primary technical achievement of this paper is a substantially faster method for computing the score function of the psmc model. First, I summarize the existing state of the art. Using the standard forward algorithm (e.g., Bishop, 2006), as in psmc, the log-likelihood of a hidden Markov model with *M* hidden states can be computed in 𝒪(*LM*^2^) time and 𝒪(*M*) storage when there are *L* sequential observations. Thus, each likelihood evaluation requires a complete pass over the data, and the algorithm must proceed serially (cannot be parallelized) because of the sequential dependency structure of the hidden Markov model. By exploiting problem-specific structure, Harris et al. (2014) improved the asymptotic running time of the psmc forward algorithm to 𝒪(*LM*), with the same storage requirement. Palamara et al. (2018) later extended this technique give 𝒪(*LM*) forward and backward algorithms for the more accurate psmc ′ model.

To numerically differentiate the log-likelihood, one obvious approach is to apply automatic differentiation to the linear-time forward algorithm of Palamara et al. This is very easy to implement using differentiable programming languages designed for neural networks (Martín Abadi et al., 2015; Bradbury et al., 2018). However, from a performance perspective, it is suboptimal. Using reverse-mode automatic differentiation, the algorithm has to store all intermediate computation values, requiring 𝒪(*LM*) bytes of storage. This results in very high memory overhead; in particular it renders the resulting procedure unsuitable for running on a graphics processing unit (GPU), which is currently the main growth area in high performance computing. Alternatively, one can employ forward-mode automatic differentiation, which works by tracking a set of “dual numbers” alongside the primal computation. However, this has high computational overhead, since every floating-point operation results in. 𝒪(*M*) additional dual operations, and it is not as optimized, since neural networks rely almost exclusively on back-propagation for training. In my initial experiments, both approaches required 10–30 seconds per gradient evaluation (Figure 4a), too slow to be useful in practice.

To achieve greater performance, I developed a novel score function algorithm which has. 𝒪(*LM* ^2^) time and 𝒪(*M* ^2^) storage complexity. In other words, the algorithm gives gradients “for free” in the same amount of time that it takes the naïve forward algorithm to evaluate the likelihood of an HMM, and its low memory requirement renders it suitable for running massively in parallel on a GPU. Numerical experiments (see Figure 4a) show that it is around 30–90x faster than automatic differentiation. In one sentence, the algorithm works by exploiting a classical identity due to R.A. Fisher for computing the score function of a latent variable model, and accelerating the computations by harnessing problem-specific structure in a manner similar to Palamara et al. (2018). The precise technical explanation requires much additional notation, and is not necessary to understand the rest of the paper, so it is deferred to Appendix A.

#### 2.3.2 Parallel evaluation

The preceding subsection shows how it is possible to evaluate the psmc score function in a memory-limited environment, such as on a GPU. However, each evaluation requires a complete pass over the chromosome, which is expensive when there are many chromosomes and the genome length is long. To achieve greater speed, I employ a further approximation which breaks up long stretches of the chromosome into smaller “chunks” that are effectively independent. The idea is to exploit the fact that a hidden Markov model forgets its initial state quickly in the sense that, for sufficiently large *f* > 0,

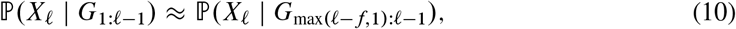

where *X*_*𝓁*_ and *G*_*1:𝓁*_ = (*G*_1_, …,*G*) are as in (1)-(3). In other words, the predictive distribution ℙ(*X*_*𝓁*_ | *G*_1: *𝓁*-1_) is effectively unchanged if, instead of conditioning on the entire observation sequence up to position *𝓁*, we instead condition only up to a fixed window before it. This is a result of the fact that the hidden Markov model forgets its initial state exponentially quickly, so that *X*_*𝓁*_ is approximately independent of *G*_1: *𝓁−f*−1_ conditional on *G*_*𝓁*−*f:𝓁*−1_ (Le Gland and Mevel, 2000).

##### Algorithm 1 Parallel log-likelihood

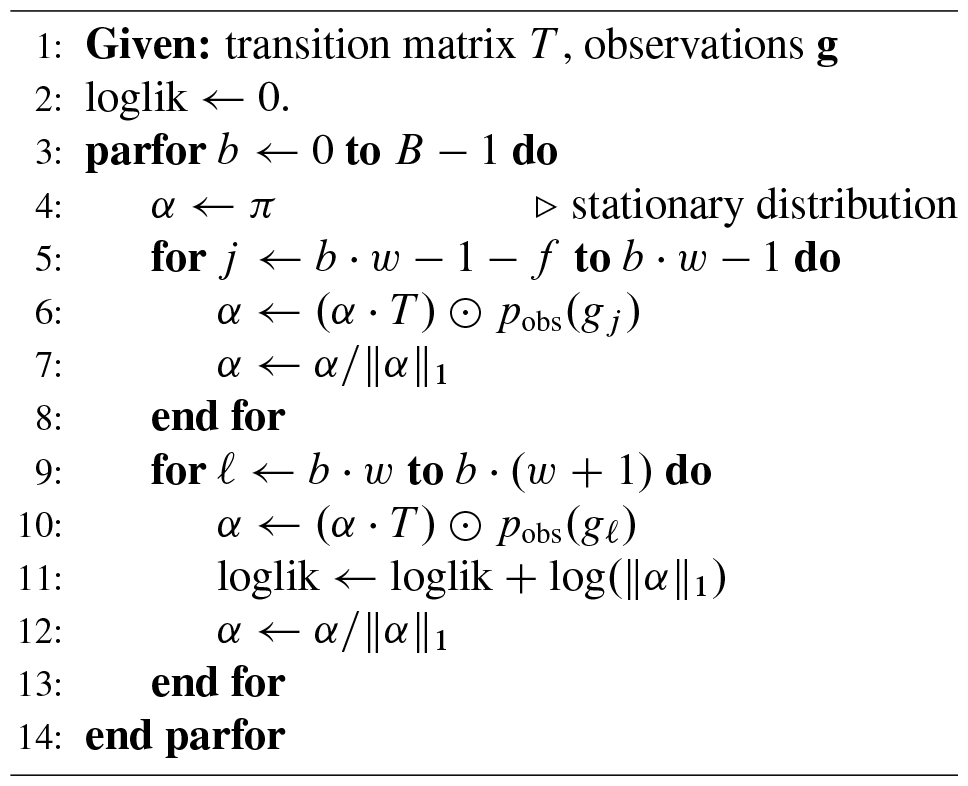

The approximation immediately leads to a parallelizable stochastic gradient algorithm for ascending the psmc likelihood surface: selecting a “window size” *w* such that *B* = *L/w* (assume for simplicity *w* evenly divides *L*),

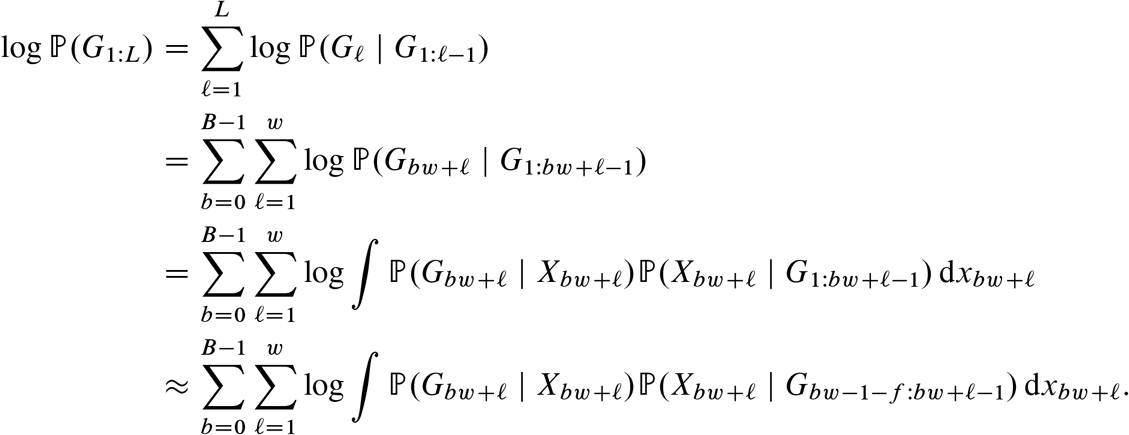

Then observe that a simple recursion, displayed in Figure 1, can be used to compute each of the “within-block” inner summations in the last line in parallel. The same idea also works for computing the score function due to the connection to the standard forward algorithm described in Appendix B. This leads to a natural stochastic gradient procedure where, instead of computing the gradient across all blocks *b* =1, *…,B* of a chromosome, we randomly sample a small number *S* « *B* of blocks, compute the score, and then return an unbiased estimator of the gradient.

**Figure 1:**
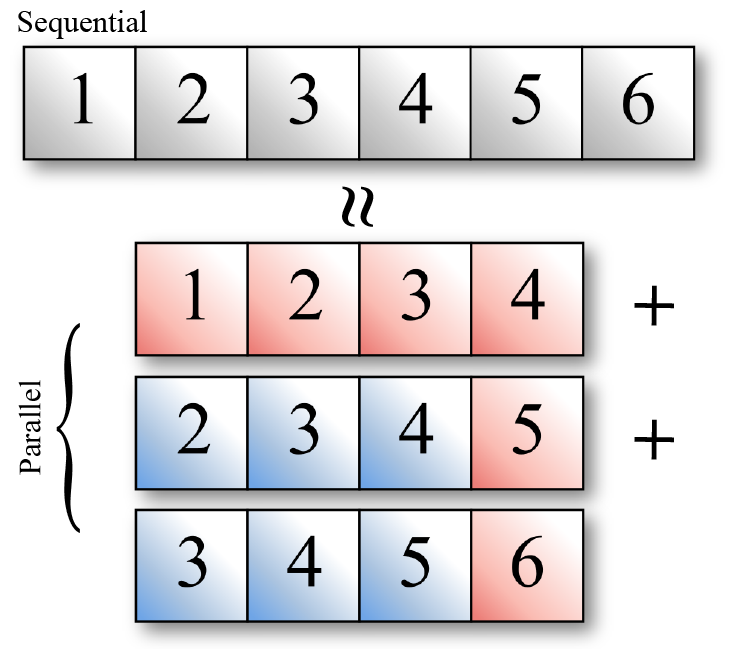
Parallel algorithm for evaluating the likelihood of an HMM. The exact algorithm evaluates the likelihood sequentially across each observation (grey blocks in figure). The parallel algorithm breaks the observation sequence up into a set of overlapping blocks which can be evaluated in parallel. Blue blocks are used to “warm up” the filtering distribution, and red blocks contribute to the likelihood calculation.

The accuracy of this approximation depends on selecting the number of blocks *B* and, particularly, the “overlap” parameter *f* which controls how many observations are used to approximate the target filtering distribution in (10). Although some exact bounds on the rate of forgetting are known (Le Gland and Mevel, 2000) which could be used to choose *f*, they depend on knowing the underlying model which is the quantity we are trying to infer in the first place. After some experimentation (see Section 3.3) I settled on *f* = 500 as providing a numerically accurate approximation to the log-likelihood across the range of simulated models considered in this paper.

### 2.4 Fitting procedure

With the fast score function estimator in hand, a variety of techniques are available for sampling from the posterior ℙ_phlash_ (*η*| **G)** A particular feature of the phlash model is that the posterior is likely to be highly multimodal: there are many low-dimensional time discretizations that can explain the data equally well. Several of the procedures mentioned above, VI in particular, are known to have difficulty fitting multimodal posterior distributions (Rezende and Mohamed, 2015). After some experimentation, I found that Stein variational gradient descent (SVGD; Liu and D. Wang, 2016) resulted in the best combination of accuracy and speed. SVGD is an optimization-based particle method which is well-suited to running on GPUs, since the particle updates can be performed in parallel. All the experiments in this paper were performed using 500 particles. Smoother approximations to the posterior can be obtained by increasing the particle count, at the expense of greater running time.

To prevent overfitting, phlash supports the ability to terminate early based on a measure of out-of-sample predictive accuracy. If supplied with an independent dataset (i.e., a held-out autosome), it will terminate if the expected log-predictive density has not increased for the past 100 iterations. All of the examples in this paper utilized this fitting procedure. For simulated data (Section 3.1), the first chromosome (in lexicographic order) was held out for each simulated dataset. For the real human data analyses (Section 4), the p arm of chromosome 1 was held out.

## 3 Results

To orient the reader, a typical output from phlash is shown in Figure 2. For most applications, the main quantities of interest will be the posterior median of the sampled size histories, shown in dark blue in the figure, as well as the corresponding credible interval, shown in light blue. In my testing I found the posterior median to have good accuracy as a point estimator of the underlying size history function; this will be systematically examined in the next section. The credible interval provides a (point-wise) estimate of uncertainty for the point estimate: it grows wider when the data are more ambiguous about effective population size at a given time point, such as for the period *t* > 10^5^ in the figure, where there are not a lot of coalescent events remaining to estimate *N*_*e*_. Finally, the credible bands delineate the smallest region such that 95% of the sampled size history functions are completely contained within the bands; they are necessarily larger than the pointwise credible intervals. The credible bands are calculated by solving an optimization problem as described in Appendix B.

**Figure 2:**
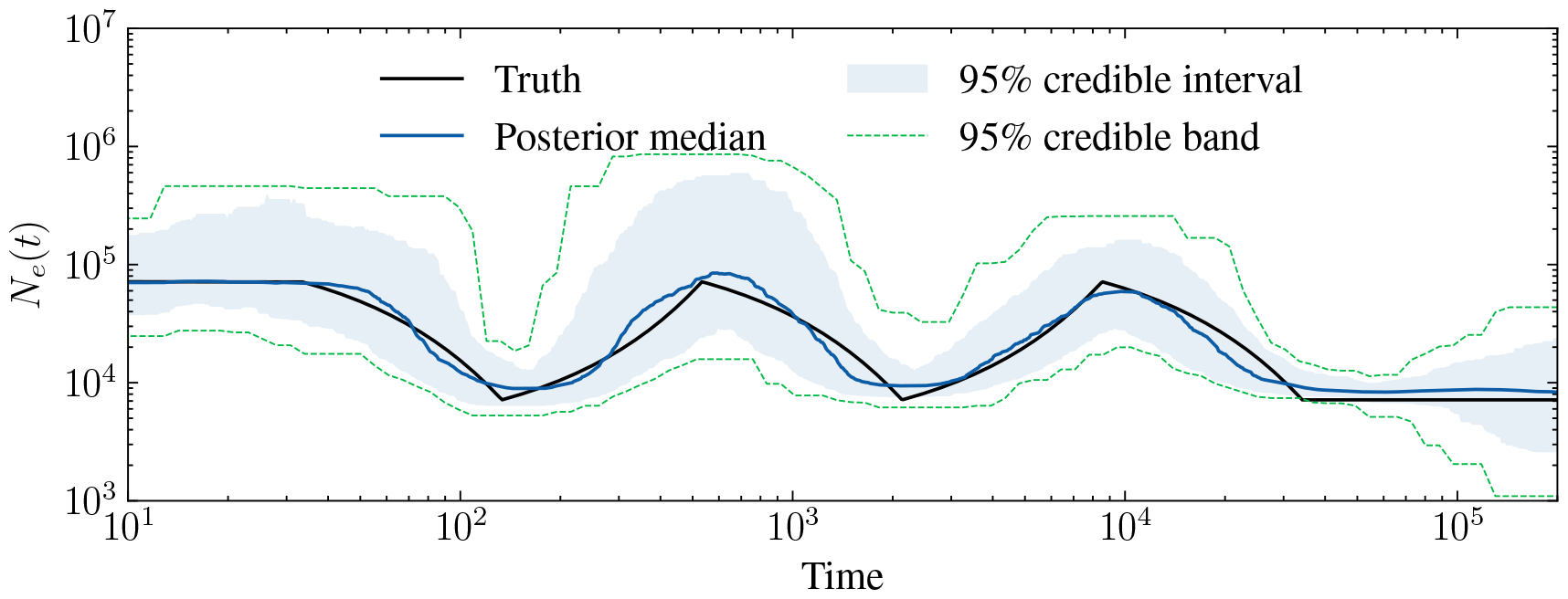
Example output of the algorithm for the “sawtooth” demography (Schiffels and Durbin, 2014; Adrion et al., 2020). *n* = 1000 diploid samples were simulated according to the size history shown in black. The posterior distribution returned by phlash is summarized by its median (dark blue line), central 95% point-wise credible interval (shaded blue region), and 95% simultaneous credible band (dashed green line).

### 3.1 Accuracy compared to existing methods

I first studied how well phlash performs on simulated data where the ground truth is known. I compared phlash to three existing methods: smc++, Msmc2, and FitCoal. smc++ (Terhorst, J. A. Kamm, and Yun S Song, 2017) is a generalization of psmc which also incorporates frequency spectrum information, by modeling the expected site frequency spectrum conditional on knowing the TMRCA of a pair of distinguished lineages. Msmc2 (Schiffels and K. Wang, 2020) optimizes a composite objective where the psmc likelihood is evaluated over all pairs of haplotypes. Finally, FitCoal (Hu et al., 2023) is a newer method which uses the site frequency spectrum to estimate size history. These three were chosen as laying at the intersection of methods that a) are relatively recent, b) have an easy-to-use software implementation, and c) can run on non-human data (i.e., do not require phased data, or detailed genetic maps).

I compared the methods across a panel of 12 different demographic models from the `stdpopsim` catalog (Adrion et al., 2020). Each model is the result of a previously published study. A particular goal of phlash is for it to be generally applicable across a wide range of biological systems. To that end, a total of eight different species are represented in the benchmark suite: *A. gambiae, A. thaliana, B. taurus, D. melanogaster, H. sapiens, P. troglodytes, P. anubis*, and *P. abelii*. Additional details of each model are shown in Table D.1. Together, these constitute a fairly diverse range of demographic models, biological parameters (i.e., mutation and recombination rates), and genome sizes.

To perform the benchmarks, I simulated whole-genome data under each of the models for diploid sample sizes *n* ∈ {1, 10, 100, 1000}. For reasons of budget and carbon footprint, only three replicates were performed for each model, resulting in a total of 12 × 4 × 3 = 144 different simulation runs. For some species in the panel, such as *A. gambiae*, the population-scaled recombination rate is so high that the default simulation engine for `stdpopsim` (msprime, Baumdicker et al., 2022), did not terminate after 24h. Thus, for uniformity, all simulations were performed using the coalescent simulator Scrm (Staab et al., 2015).

I ran each inference method on each of the simulated datasets. All methods were limited to 24h of wall time and 256Gb of RAM. This meant that it was only possible to run some of the methods for certain sample sizes: smc++ could only analyze *n* ∈{1, 10} in the allotted amount of time; Msmc2 could only analyze *n* ∈{1, 10} in the allotted amount of memory, and FitCoal could only analyze *n* ∈{10, 100} in the allotted amount of time (it crashed with an error for *n* = 1). The command lines used for simulation and estimation are listed in Appendix C, and summarized plots of every model fit are provided in Figures D.1–D.4.

I considered three accuracy metrics. The first, root mean-squared (or *L*_2_) error, has previously been used to compare the accuracy of demographic inference methods (e.g., Robinson et al., 2014; Sellinger, Abu-Awad, and Tellier, 2021). It is defined as

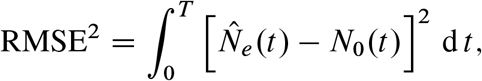

where *N*_0_(*t*) is the true historical effective population size that was used to simulate data, and *T* is an arbitrarily-chosen time cutoff (I chose *T* = 10^6^ generations). Overall results are shown in Table 1. No method uniformly dominates, but overall, phlash tended to be the most accurate most often, and is at least competitive with the more accurate method when it is not: of the 4 × 12 = 48 scenarios considered, phlash was the most accurate *30=48* D *62:5*% of the time, compared with *7=48* for FitCoal, *6=48* for smc++, and *5=48* for Msmc2. For the *n* = 1 scenarios where only a single diploid genome is available, the difference in performance between phlash and smc++/Msmc2 tended to be small, and sometimes in favor of the latter methods. I attribute this to the non-parametric nature of the phlash estimator, which does not benefit from as much prior regularization and generally needs more data to perform well. Also, as can be seen for the `Constant` model, FitCoal is extremely accurate when the true underlying model is a member of its assumed model class, i.e. containing a small number of epochs of constant or exponential growth. However, it is not clear that this is a reasonable assumption for natural populations; indeed the real data results presented below seem to suggest otherwise.

**Table 1:** Root mean-squared error for simulated data. All results have been divided by 10^8^ for clarity. Each entry is averaged over three simulation replicates. **Bold** marks the entry with the lowest mean in each row. Entries with * are significantly lower than all other entries in the same row using a Bonferroni-corrected *t* -test with FWER = 0.05.

| Model | $n$ | SMC++ | MSMC2 | PHLASH | FITCOAL |
| --- | --- | --- | --- | --- | --- |
| Africa_1T12 | 1 | 0.0232 | 0.0352 | **0.0184\*** | — |
|  | 10 | 0.0242 | 0.622 | **0.0209** | 0.0238 |
|  | 100 | — | — | **0.0166\*** | 0.0238 |
|  | 1000 | — | — | **0.0134** | — |
| African3Epoch_1H18 | 1 | 0.2 | **0.179** | 1.28 | — |
|  | 10 | 0.258 | 0.235 | **0.198** | 0.237 |
|  | 100 | — | — | 0.148 | **0.0767\*** |
|  | 1000 | — | — | **0.147** | — |
| African3Epoch_1S16 | 1 | **1.8** | 3.64 | 2.5 | — |
|  | 10 | 1.52 | 4.21 | 0.635 | **0.273\*** |
|  | 100 | — | — | 0.548 | **0.232\*** |
|  | 1000 | — | — | **0.68** | — |
| AmericanAdmixture_4B11 | 1 | **0.0134** | 0.988 | 4.66 | — |
|  | 10 | **0.0188** | 0.0569 | 0.126 | 0.0222 |
|  | 100 | — | — | 0.0723 | **0.0283\*** |
|  | 1000 | — | — | **0.0487** | — |
| BonoboGhost_4K19 | 1 | 0.374 | **0.289\*** | 0.324 | — |
|  | 10 | 0.375 | **0.289\*** | 0.357 | 0.358 |
|  | 100 | — | — | 0.378 | **0.358** |
|  | 1000 | — | — | **0.392** | — |
| Constant | 1 | **0.00579** | 0.0428 | 0.0102 | — |
|  | 10 | 0.00597 | 0.0145 | 0.00692 | **0.00009\*** |
|  | 100 | — | — | 0.00732 | **0.000123\*** |
|  | 1000 | — | — | **0.00359** | — |
| GabonAg1000G_1A17 | 1 | 6.57 | **2.77** | 3.88 | — |
|  | 10 | 5.3 | 9.29 | **0.836** | 0.955 |
|  | 100 | — | — | **0.847\*** | 0.977 |
|  | 1000 | — | — | **0.74** | — |
| HolsteinFriesian_1M13 | 1 | 0.055 | 0.086 | **0.0503** | — |
|  | 10 | 0.0563 | 0.0553 | **0.0438** | 0.0715 |
|  | 100 | — | — | **0.0564\*** | 0.117 |
|  | 1000 | — | — | **0.0617** | — |
| SinglePopSMCpp_1W22 | 1 | 0.208 | 0.547 | **0.123** | — |
|  | 10 | 0.198 | 0.121 | **0.108\*** | 0.167 |
|  | 100 | — | — | **0.109\*** | 0.161 |
|  | 1000 | — | — | **0.11** | — |
| SouthMiddleAtlas_1D17 | 1 | 0.241 | **0.152** | 0.157 | — |
|  | 10 | 0.237 | 0.164 | **0.163** | 0.555 |
|  | 100 | — | — | **0.127\*** | 0.388 |
|  | 1000 | — | — | **0.174** | — |
| TwoSpecies_2L11 | 1 | **0.0122** | 0.044 | 0.0154 | — |
|  | 10 | 0.0114 | 0.0184 | **0.0112** | 0.154 |
|  | 100 | — | — | **0.0176\*** | 0.0862 |
|  | 1000 | — | — | **0.0128** | — |
| Zigzag_1S14 | 1 | **0.0265** | 0.0642 | 0.157 | — |
|  | 10 | 0.0337 | 0.449 | **0.0118** | 0.0235 |
|  | 100 | — | — | **0.0132\*** | 0.0324 |
|  | 1000 | — | — | **0.014** | — |

Although RMSE is a natural error measure, it does not necessarily paint a complete picture, since portions of the size history function may be effectively inestimable by any method due to the coalescent sampling process. This can happen, for example, when estimating very recent population history in an exponentially growing population, or very ancient history in a population that has undergone a bottleneck. In both cases, lack of coalescent events render these time periods “invisible” to coalescent-based demographic inference methods (Terhorst and Yun S Song, 2015; Baharian and Gravel, 2018). In these time periods, *L*_2_ error represents algorithm bias only since there is minimal signal to guide the estimator. An example of this phenomenon shown in Figure 3: in panel (a), phlash has very large *L*_2_ error compare to smc++ and Msmc2, because the latter two methods assume that *η* is flat until 10^4^–10^5^ generations ago, which happens to be true for this particular demographic model. However, since this model has a large *N*_*e*_ until 10^5^ generations ago, the probability of coalescence during this period is low (Figure 3b).

**Figure 3:**
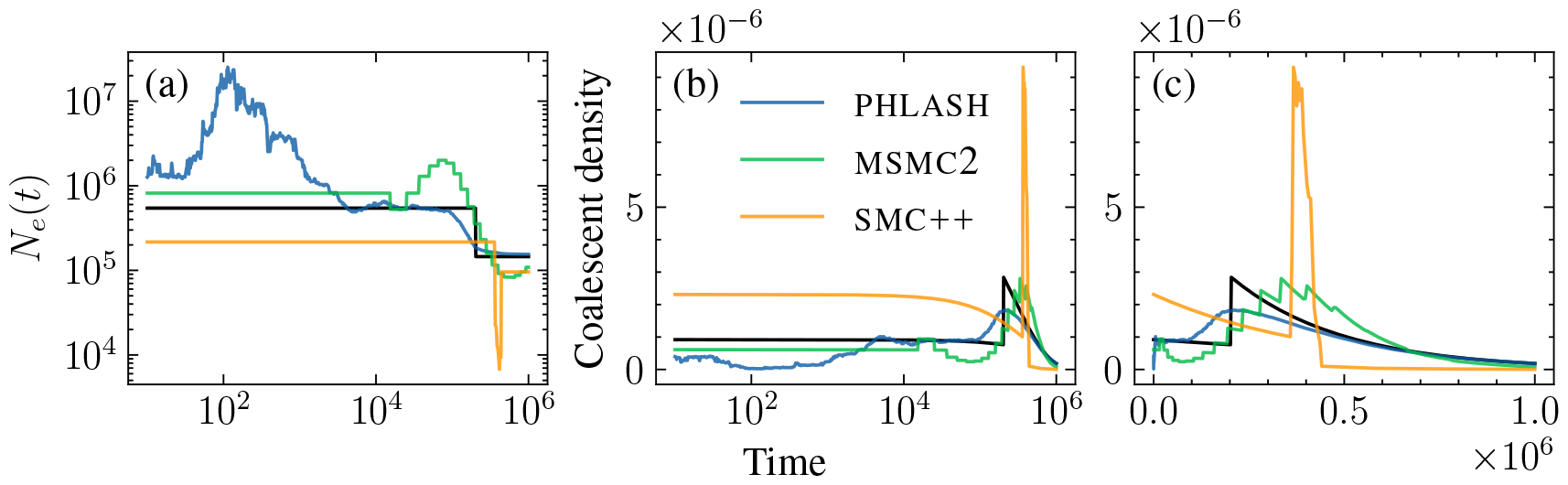
*L*_2_ versus total variation error for the `African3Epoch_S16` demography when *n* = 1. phlash has large mean-squared error relative to Msmc2 and smc++ (a), but it mainly occurs at very recent times, when there is low density of coalescence. Total variation error (b–c) accounts for this by measuring the *L*_1_ distance between the true and inferred density functions.

Since, ultimately, size history inference is a density estimation problem, a natural alternative measure is the total variation distance between inferred and true demographic models. This is defined as

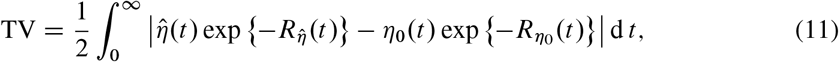

where 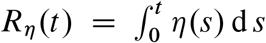 is the cumulative hazard rate of coalescence up to time *t*, 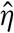 is the estimate, and *η*_0_ is the truth. An example of the integrands is shown in Figure 3(c); phlash in fact has lowest TV error for this example, despite possessing the largest mean-squared error.

Finally, I considered an additional error metric derived from the fact that the coalescent intensity function *η* actually indexes an entire *family* of distributions, since it also gives the time to first coalescence in a sample of size *n*, with hazard rate of coalescence 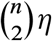. Define the “scaled” total variation distance

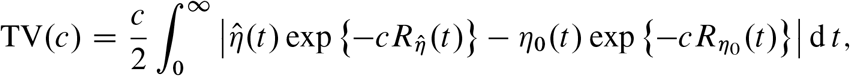

and then

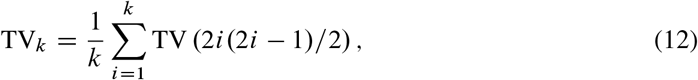

which is total variation distance 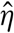 and *η*_0_ averaged over their implied first-coalescence distributions, in diploid samples of size *i* = 1,*…,k:* Note that TV(1) = TV_1_ = TV as defined in equation (11). The idea of TV_*k*_ is to measure how increasing sample sizes improves estimation accuracy in the recent past. In the results below I chose *k* = *n*, where *n* is the simulated sample size, though this choice is somewhat arbitrary.

Results for TV and TV_*n*_ are shown in Tables 2 and 3. When considering TV error, phlash was most accurate two-thirds of the time (32 of 48 scenarios), and 75% of the time (36/48) when considering TV_*n*_. A few other aspects are worth considering. It can be seen that TV error does not always decrease with increasing sample size *n* = 1 →10 → 100 → 1000, though it does tend to shrink the confidence intervals. Some possible causes and solutions are discussed in Section 5. This was also observed for TV_*n*_, though we should not necessarily expect it to decrease with *n*, since by (12), it places greater emphasis on accuracy in the very recent past as *n* grows. Finally, it must be noted many of the models in the `stdpopsim` catalog were in fact estimated using a psmc-type model, so in that sense the catalog is enriched for members of the simplified model spaces considered by these programs. Hence there is liable to be a small, intangible bias against nonparametric methods in these benchmarks.

**Table 2:** Total-variation error for simulated data. Formatting is same as Table 1.

| Model | $n$ | SMC++ | MSMC2 | PHLASH | FITCOAL |
| --- | --- | --- | --- | --- | --- |
| Africa_1T12 | 1 | 0.0887 | 0.052 | **0.0408\*** | — |
|  | 10 | 0.107 | 0.0476 | **0.0375\*** | 0.066 |
|  | 100 | — | — | **0.0369\*** | 0.0661 |
|  | 1000 | — | — | **0.0257** | — |
| African3Epoch_1H18 | 1 | 0.113 | 0.0874 | **0.0832** | — |
|  | 10 | 0.139 | **0.0578\*** | 0.065 | 0.102 |
|  | 100 | — | — | 0.0514 | **0.0306** |
|  | 1000 | — | — | **0.0448** | — |
| African3Epoch_1S16 | 1 | 0.548 | 0.21 | **0.0886\*** | — |
|  | 10 | 0.563 | 0.199 | 0.0545 | **0.0132\*** |
|  | 100 | — | — | 0.0452 | **0.0106\*** |
|  | 1000 | — | — | **0.063** | — |
| AmericanAdmixture_4B11 | 1 | 0.121 | **0.0743** | 0.0747 | — |
|  | 10 | 0.165 | **0.0649\*** | 0.118 | 0.188 |
|  | 100 | — | — | **0.151\*** | 0.196 |
|  | 1000 | — | — | **0.153** | — |
| BonoboGhost_4K19 | 1 | 0.307 | 0.159 | **0.152** | — |
|  | 10 | 0.279 | 0.155 | **0.111\*** | 0.629 |
|  | 100 | — | — | **0.139\*** | 0.239 |
|  | 1000 | — | — | **0.16** | — |
| Constant | 1 | 0.0533 | 0.0245 | **0.0232** | — |
|  | 10 | 0.0549 | 0.0218 | 0.0179 | **0.000319\*** |
|  | 100 | — | — | 0.0159 | **0.000454\*** |
|  | 1000 | — | — | **0.0122** | — |
| GabonAg1000G_1A17 | 1 | 0.527 | 0.157 | **0.0423\*** | — |
|  | 10 | 0.524 | 0.156 | 0.0267 | **0.00424** |
|  | 100 | — | — | 0.0326 | **0.00452\*** |
|  | 1000 | — | — | **0.0277** | — |
| HolsteinFriesian_1M13 | 1 | 0.15 | 0.125 | **0.0922\*** | — |
|  | 10 | 0.185 | **0.096** | 0.0977 | 0.39 |
|  | 100 | — | — | **0.0883\*** | 0.444 |
|  | 1000 | — | — | **0.0867** | — |
| SinglePopSMCpp_1W22 | 1 | 0.245 | 0.0953 | **0.0807** | — |
|  | 10 | 0.249 | 0.0935 | **0.0801** | 0.113 |
|  | 100 | — | — | **0.0798** | 0.0961 |
|  | 1000 | — | — | **0.0803** | — |
| SouthMiddleAtlas_1D17 | 1 | 0.0572 | 0.0491 | **0.0487** | — |
|  | 10 | 0.0621 | **0.0425** | 0.0504 | 0.108 |
|  | 100 | — | — | **0.0365\*** | 0.0816 |
|  | 1000 | — | — | **0.0423** | — |
| TwoSpecies_2L11 | 1 | 0.0267 | **0.0147\*** | 0.0314 | — |
|  | 10 | 0.0249 | **0.0132** | 0.0342 | 0.172 |
|  | 100 | — | — | **0.0296\*** | 0.138 |
|  | 1000 | — | — | **0.0224** | — |
| Zigzag_1S14 | 1 | 0.142 | **0.0666\*** | 0.0899 | — |
|  | 10 | 0.136 | **0.0667\*** | 0.0862 | 0.103 |
|  | 100 | — | — | **0.0688\*** | 0.143 |
|  | 1000 | — | — | **0.0695** | — |

**Table 3:** TV_*n*_ error for simulated data. See main text for description of this metric. Formatting is same as Table 1.

| Model | $n$ | SMC++ | MSMC2 | PHLASH | FITCOAL |
| --- | --- | --- | --- | --- | --- |
| Africa_1T12 | 1 | 0.0887 | 0.052 | **0.0408\*** | — |
|  | 10 | 0.245 | **0.0965\*** | 0.109 | 0.44 |
|  | 100 | — | — | **0.268\*** | 0.856 |
|  | 1000 | — | — | **0.138** | — |
| African3Epoch_1H18 | 1 | 0.113 | 0.0874 | **0.0832** | — |
|  | 10 | 0.264 | 0.228 | 0.216 | **0.214** |
|  | 100 | — | — | **0.0572** | 0.0836 |
|  | 1000 | — | — | **0.0291** | — |
| African3Epoch_1S16 | 1 | 0.548 | 0.21 | **0.0886\*** | — |
|  | 10 | 0.124 | 0.229 | **0.0849\*** | 0.22 |
|  | 100 | — | — | **0.042\*** | 0.429 |
|  | 1000 | — | — | **0.0271** | — |
| AmericanAdmixture_4B11 | 1 | 0.121 | **0.0743** | 0.0747 | — |
|  | 10 | 0.318 | **0.145\*** | 0.542 | 0.468 |
|  | 100 | — | — | **0.156\*** | 0.631 |
|  | 1000 | — | — | **0.045** | — |
| BonoboGhost_4K19 | 1 | 0.307 | 0.159 | **0.152** | — |
|  | 10 | 0.136 | 0.124 | **0.0715\*** | 0.193 |
|  | 100 | — | — | **0.0292\*** | 0.441 |
|  | 1000 | — | — | **0.0196** | — |
| Constant | 1 | 0.0533 | 0.0245 | **0.0232** | — |
|  | 10 | 0.0792 | 0.0302 | 0.0289 | **0.000319\*** |
|  | 100 | — | — | 0.0137 | **0.000454\*** |
|  | 1000 | — | — | **0.0257** | — |
| GabonAg1000G_1A17 | 1 | 0.527 | 0.157 | **0.0423\*** | — |
|  | 10 | 0.36 | 0.405 | 0.171 | **0.167\*** |
|  | 100 | — | — | **0.309\*** | 0.363 |
|  | 1000 | — | — | **0.457** | — |
| HolsteinFriesian_1M13 | 1 | 0.15 | 0.125 | **0.0922\*** | — |
|  | 10 | **0.453** | 0.643 | 0.514 | 0.89 |
|  | 100 | — | — | **0.497\*** | 0.978 |
|  | 1000 | — | — | **0.507** | — |
| SinglePopSMCpp_1W22 | 1 | 0.245 | 0.0953 | **0.0807** | — |
|  | 10 | 0.207 | 0.152 | **0.0843\*** | 0.185 |
|  | 100 | — | — | **0.204\*** | 0.422 |
|  | 1000 | — | — | **0.0115** | — |
| SouthMiddleAtlas_1D17 | 1 | 0.0572 | 0.0491 | **0.0487** | — |
|  | 10 | **0.0911** | 0.139 | 0.118 | 0.243 |
|  | 100 | — | — | **0.0261\*** | 0.315 |
|  | 1000 | — | — | **0.0183** | — |
| TwoSpecies_2L11 | 1 | 0.0267 | **0.0147\*** | 0.0314 | — |
|  | 10 | 0.0616 | **0.0307** | 0.0948 | 0.415 |
|  | 100 | — | — | **0.0487\*** | 0.377 |
|  | 1000 | — | — | **0.104** | — |
| Zigzag_1S14 | 1 | 0.142 | **0.0666\*** | 0.0899 | — |
|  | 10 | 0.278 | 0.22 | **0.175\*** | 0.393 |
|  | 100 | — | — | **0.105\*** | 0.674 |
|  | 1000 | — | — | **0.00889** | — |

### 3.2 Running time and memory consumption

Next, I examined the computational resources required by each method when analyzing the simulated datasets mentioned above. The peak amount of memory used, as well as total CPU time, was recorded for each simulation run. Because the datasets are simulated from organisms with differing genome lengths, I normalized both of these measures by genome length (measured in gigabase-pairs) to make them comparable across runs, and then averaged the data from all runs together for each method and sample size.

The results are shown in Figure 4(b)-(c). For analyzing a single diploid sample, *n* = 1, all methods required a similar amount of CPU time, around 20-30 minutes per gigabase; Msmc2 required the most memory, and phlash required the least. For *n* = 10, the only sample size where it was possible to run all four methods, FitCoal was the most efficient in terms of time and memory usage, which was expected since it only analyzes the frequency spectrum. Of the HMMbased approaches, phlash required significantly less CPU time and memory than smc++ and, especially, Msmc2. Increasing the sample size to *n* = 100 caused the running time of FitCoal to increase by roughly tenfold, while memory consumption remained low, while for phlash, CPU and memory demands increased only moderately. Finally, for *n* = 1000, no method except phlash was able to run given the allotted computational resources, and one thousand diploid samples using phlash required roughly the same amount of memory, and less CPU time, than analyzing ten samples using Msmc2.

**Figure 4:**
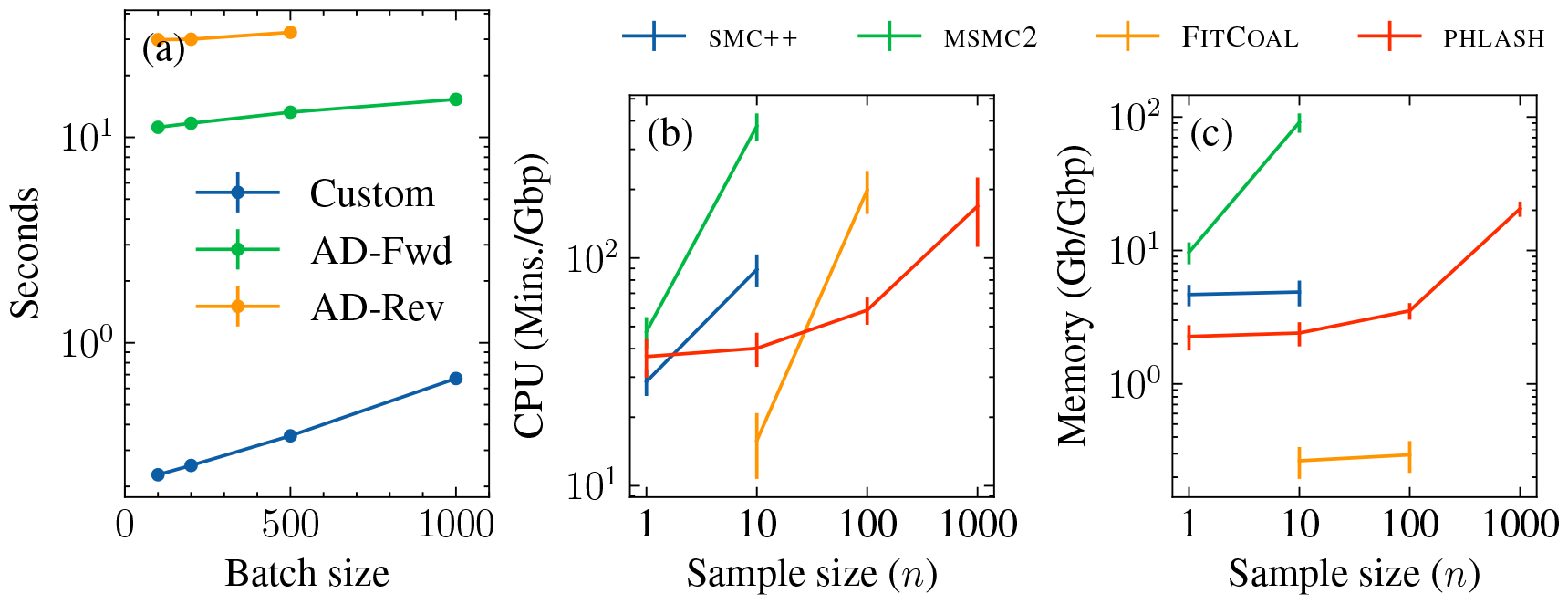
Benchmarking results. (a) Time needed to evaluate score function as a function of batch size. The sequence length was *L* = 10^5^. Custom algorithm is compared forward- and reverse-mode automatic differentiation of the accelerated/linear-time forward algorithm using Jax (Bradbury et al., 2018). For *n* = 1000, reverse-mode automatic differentiation exhausted the available GPU memory and was unable to be evaluated. (b) Mean CPU time and (c) peak memory usage for the various methods, with standard error bars.

### 3.3 Accuracy of the parallel approximation

As mentioned above, the key feature that enables phlash to analyze large datasets quickly is that it runs in parallel on many GPU cores. Although it is somewhat implied by the simulation results of the preceding section, I directly verified that this approximation is accurate and does not introduce too much numerical error into the computations. Specifically, for each of the simulation runs, I compared the results of a) evaluating the log-likelihood using the exact (i.e., sequential) forward algorithm and b) the parallel algorithm, for different settings of the overlap parameter *f* (cf. Section 2.3.2). Smaller values of *f* lead to greater computational gains, since there is less sequence overlap, but more approximation error, since the Markov chain does not have as much time to forget its initial distribution.

Results are shown in Figure 5(a). When *f* = 0, the method simply breaks the chromosome into non-overlapping chunks and analyzes them in parallel; as expected, relative error is highest at this setting. Increasing the overlap length from *f* = 100 up to *f* = 1000 successively diminished the numerical error. At *f* = 500 the relative error is about 10^−8^, roughly the smallest difference that can be represented using single-precision floating point. Since the GPU implementation also runs in single precision, I chose *f* = 500 for all the results reported in the paper. Finally, I note the presence of a few outliers in Figure 5(a), which correspond to organisms and demographic models where the approximation error diminishes less rapidly than average. Generally this occurs in inbred populations, where long runs of homozygosity appear to lessen the rate of forgetting in the coalescent Markov chain. Indeed, the model with the highest relative error was `HolsteinFriesian_1M13`, which features a very recent population crash. For species with similar characteristics, users may want to consider increasing *f*.

**Figure 5:**
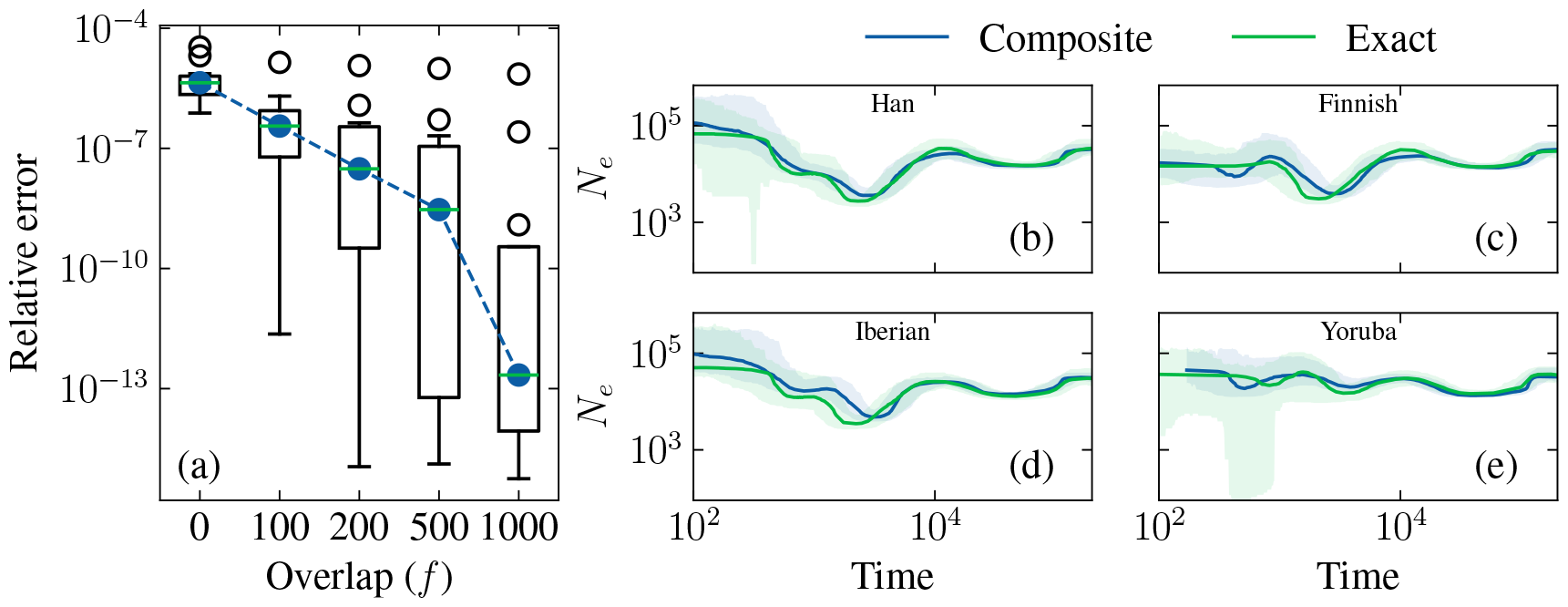
Accuracy of likelihood approximations. Panel (a) is a box-whisker plot of the relative error, defined as |loglik_exact_ − loglik_parallel_| /loglik_exact_, for various settings of the overlap parameter *f*, across all simulations. Panels (b)–(e) compare the results of model fitting using the composite likelihood approximate, versus independent likelihoods (see text for description).

### 3.4 Calibration of the composite likelihood posterior

Finally, I studied the extent to which the composite likelihood approximation used in phlash affects the resulting posterior distribution. In the frequentist setting, it is well-known how to adjust the Fisher information matrix in order to obtain asymptotically valid confidence intervals for the maximum composite likelihood estimator. However, the theory of Bayesian inference using composite likelihood is still in its early stages, and it is not as obvious how to adjust the posterior distribution of a composite Bayesian model in order to ensure proper calibration (Pauli, Racugno, and Ventura, 2011; Ribatet, Cooley, and Davison, 2012).

To assess the extent to which the credible intervals in phlash are deflated, I compared the estimation results on several human real data examples under two scenarios. The first scenario, “composite”, is the default setting for phlash and utilizes both frequency spectrum and linkage information as described in Section 2.2. For the second scenario, “exact”, I separated the data into two independent and roughly equally-sized subsets consisting of chromosomes 1–8 and 9–22. On the first eight chromosomes, I extracted data from a single diploid member of the population, and for the remaining ones, I estimated a site frequency spectrum at approximately unlinked sites by considering only 100bp segments at regular intervals of 25kbp. For model fitting, the psmc and SFS likelihoods were evaluated separately on these respective sources of information, leading to a model where the two likelihood terms are approximately independent of each other, and the constituent entries of the SFS are also approximately independent.

The results of these analyses are shown in Figure 5(b)–(e). There is fairly good agreement between the two scenarios, particularly in the ancient past. The “composite” scenario tends to produce narrower confidence intervals in the very recent past, though the differences do not appear extreme. There is particularly good agreement between the two sets of estimates in the ancient past, a fact which will be revisited in Section 4.4. In general, phlash estimates using the default model seem to be relatively well calibrated, and users who desire stricter calibration can employ the downsampling scheme outlined above.

### 3.5 Inferring recombination rates

phlash returns a full posterior distribution over recombination rate. Although it is not the main focus, I examined the accuracy of this posterior distribution for each of the simulated datasets described above. Results are shown in Figure 6. For reasons that are currently unclear to me, there is a slight downward bias across all models. However, in general, the estimates are fairly accurate, with a posterior median that is within 5-15% of the ground truth. Increasing the sample size sharpens the posterior distribution to an extent, with the greatest jump occurring at *n* = *1→n* = 10. Accuracy was worse for models 6 and, especially, 10; again I do not currently understand the cause, though it does not appear to adversely impact estimation accuracy (cf. Table 1 and Figures D.1 and D.4). Finally, I note that phlash currently assumes a single, uniform recombination rate across all contigs, whereas the rate is chromosome-specific in the `stdpopsim` catalog. Allowing the rate to vary by chromosome could be an avenue for future work.

**Figure 6:**
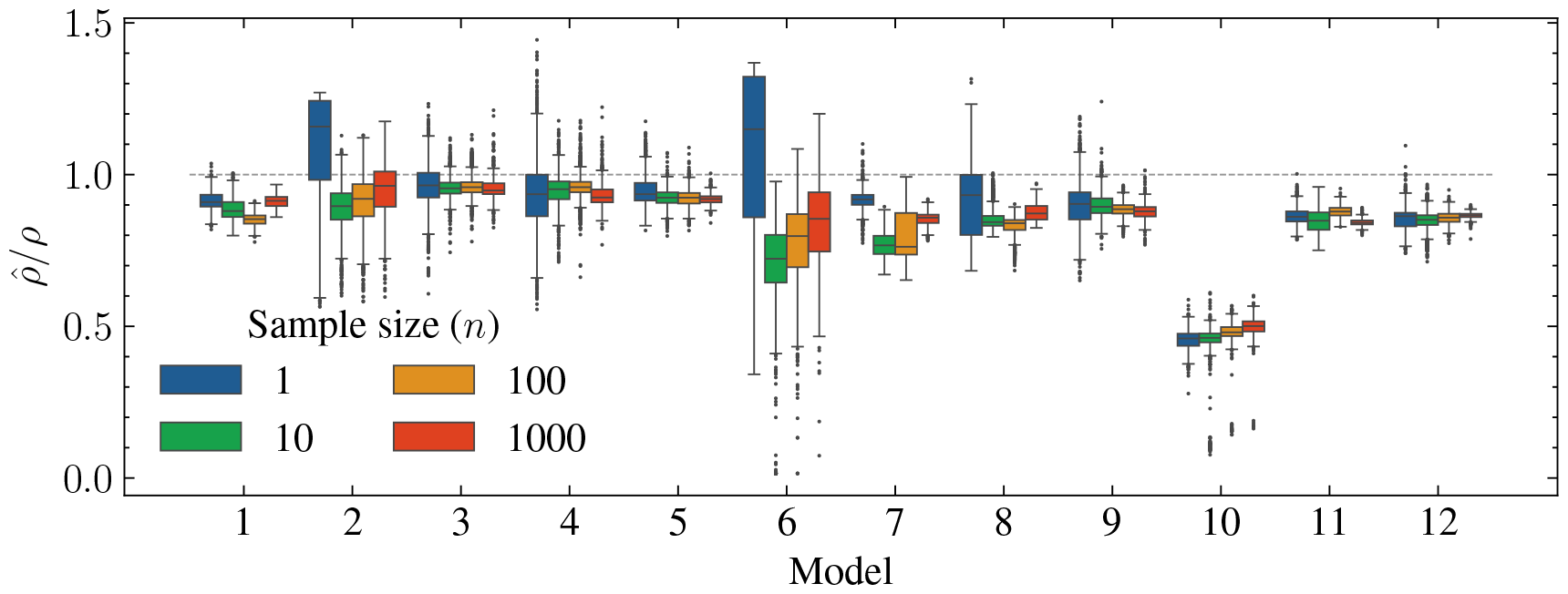
Posterior distribution of the recombination rate for each of the models shown in Table D.1.

## 4 Applications

Having verified that phlash performed well on simulated examples, I turned to analyzing real data. The analyses in this section are based on a recently published unified genomes dataset generated by Wohns et al. (2022). The dataset contains 3,601 modern samples, as well as several ancient samples, obtained from a variety of sources (Meyer et al., 2012; Prüfer et al., 2014; The 1000 Genomes Project Consortium, 2015; Mallick et al., 2016; Bergström, McCarthy, et al., 2020; Mafessoni et al., 2020). The samples are organized into 214 different subpopulations, however there is some duplication amongst them—for example, HGDP, SGDP, and 1000 Genomes all contain samples from the Yoruba population. After merging samples from duplicated populations, 159 populations remained; all of the descriptors below refer to the data after merging. Except where noted, all of the estimates assume a human mutation rate of 1:29×10^−8^ per base pair per generation and a generation time of *g* = 29 years.

### 4.1 Inferring demographic history

The main use of phlash is to infer population history. Figure 7(a) shows the results of running phlash on all 159 populations in the dataset: each opaqued black line is the posterior median for one of the populations. For clarity, I also grouped the populations into six geographical superpopulations, plus one for the ancient samples, and ran phlash on the combined data (Figure 7b). The estimates recover some of the known motifs of human evolution, including shared ancestry in the ancient past, the African/non-African divergence, recent rapid expansion, a deep divergence between the African and non-African lineages, divergence of the Oceania super-population from a Eurasian ancestral population (Wollstein et al., 2010), and the gradual extinction of all archaic populations.

**Figure 7:**
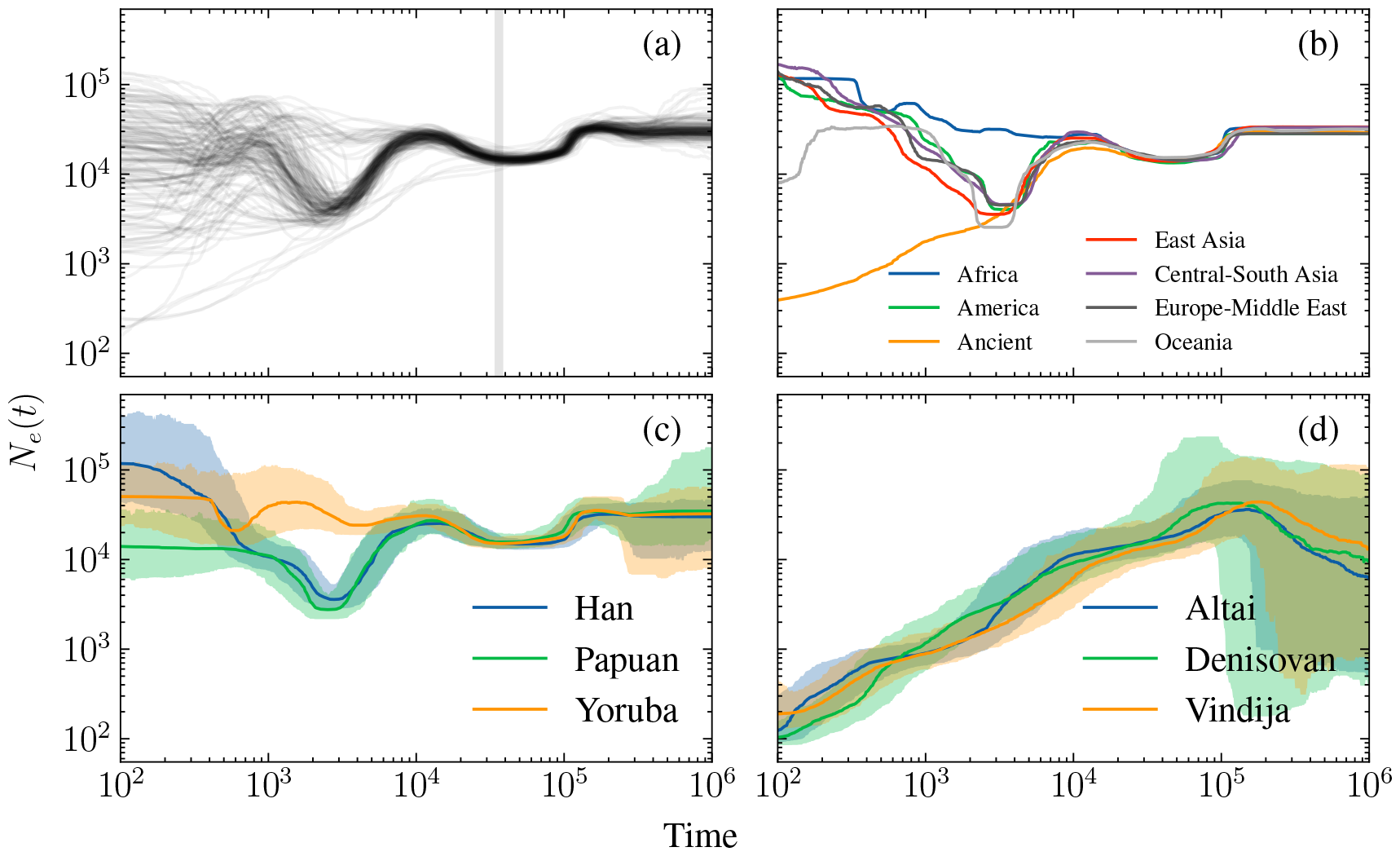
Results of running phlash on 3,609 genomes. Panel (a): estimates for all populations. Vertical shaded region is 813-930kya, see Section 4.4. (b) Estimates after grouping by super-population. (c-d): Estimates and 95% credible intervals for three (c) modern and (d) archaic subpopulations.

phlash outputs more than just point estimates: Figure 7(c) visualizes the complete posterior distribution for the Han, Yoruba, and Papuan subpopulations, as does Figure 7(d) for several archaic genomes. The width of the confidence bands can address more nuanced questions about how the populations evolved—they show, for example, increased uncertainty in the ancient as well as very recent past. Another interesting feature concerns population divergence. Panel (a) shows the African/non-African divergence occurring approximately 6,500 generations ago, which is earlier than most published estimates (e.g., López, Van Dorp, and Hellenthal, 2015). However, panel (b) reminds us that the estimates are noisy; a reasonable lower bound on the divergence time could be when the Yoruba and Han/Papuan credible bands diverge, which results in a more reasonable estimate of ∼87kya (∼3k generations ago, assuming a generation time of 29 years).

### 4.2 Detecting admixture and population structure

The preceding observation suggests a non-parametric technique for detecting population structure in pairs of populations: look for the earliest time when the effective population sizes are significantly different from each other. Formally, given two size history functions *η*_1_ and *η*_2_, a plot of the “ratio estimator” *η*_1_*/*η**_2_ + *η*_2_*=/*η**,_1_ over time should roughly reveal when the two populations diverged, assuming of course that they experienced distinct changes in effective population size after the divergence. This estimator bears some resemblance to the so-called “cross-coalescent rate” (CCR) estimator, defined by Schiffels and Durbin (2014) as the ratio *2*η**_12_*/(*η**_1_ + *η*_2_), where *η*_12_ denotes the instantaneous rate of coalescence between a haplotype sampled from population 1 and one sampled from population 2. CCR curves have been proposed as a non-parametric method of detecting population divergence and admixture (Spence et al., 2018; Schiffels and K. Wang, 2020). However, there is one important difference: estimating *η*_12_ requires accurately phased data, which may be difficult to obtain in practice, whereas the ratio estimator does not.

To explore this idea further, I simulated data for the Yoruba, Han, and European populations according to a published out-of-Africa model (Gutenkunst et al., 2009), and computed the CCR and ratio curves. Then, I estimated the same sets of curves using real data for these three populations. Results are shown in Figure 8. (Although CCR is usually constrained to lie in [*0; 1*] by jointly estimating *η*_1_, *η*_2_, and *η*_12_, phlash currently lacks this capability, so the CCR curves shown are based on independent estimates and can exceed 1.) In this model, basal CEU/CHB population diverges from YRI around 6,500 generations ago, with residual gene flow until the present. The shaded region in panels (a) and (b) highlights the period between the YRI/basal divergence, and the divergence of CEU and CHB. In both the real and simulated data, the ratio estimator has a pronounced spike around this time, which simply reflects the fact that the divergent population experienced a population crash immediately after the split. The CCR also shows an the expected transition from CCR ≈ 1 to CCR « 1 around this time, however it is perhaps a bit harder to discern. Overall, both estimators can be useful in practice. When a large/ancient split has occurred, the ratio estimator can be used to date it with decent accuracy, while the CCR curve will be more sensitive when the data can be confidently phased.

**Figure 8:**
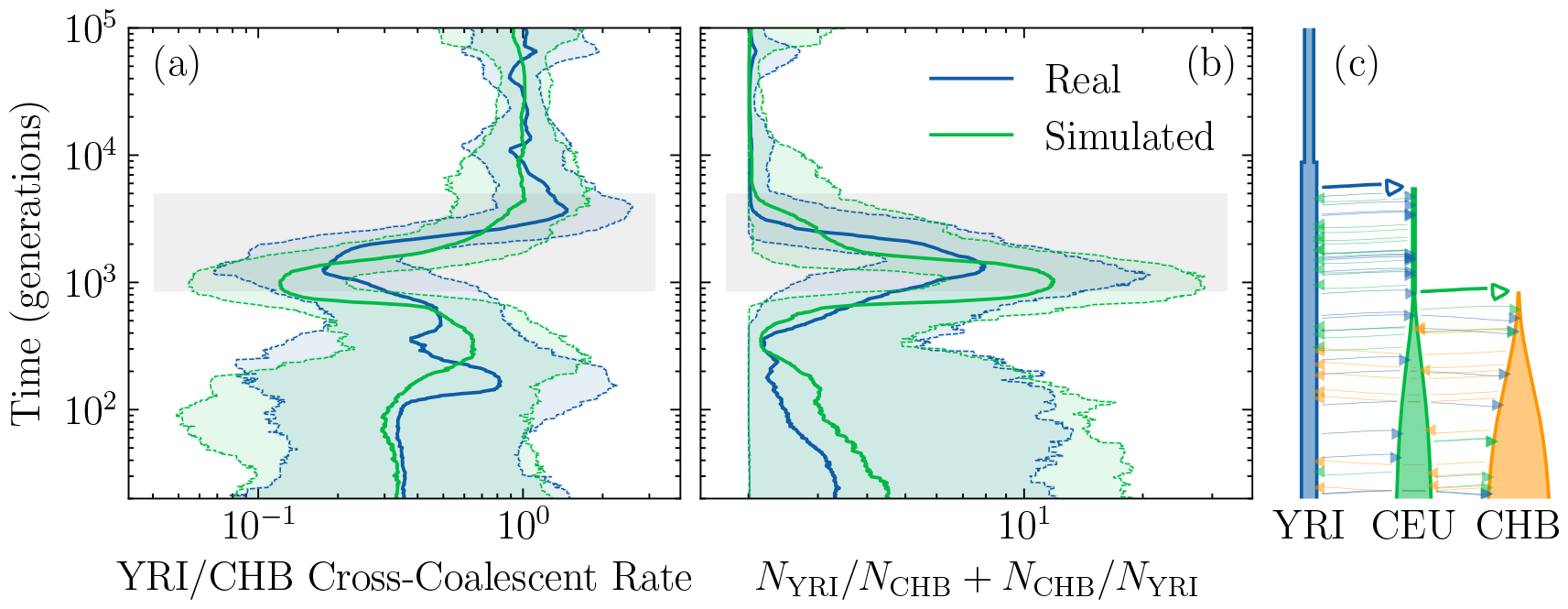
Posterior inference of population structure. (a) Posterior distribution (median + 95% credible interval) of cross-coalescent rate between the Yoruba (YRI) and Han (CHB) populations. (b) Posterior distribution of non-parametric ratio estimator. (c) Three-population out-of-Africa model used to generate simulated data.

### 4.3 Analyzing ancient DNA

phlash can be used on ancient samples without modification. In this case, the time units should be interpreted as “generations before the lifetime of the sample” instead of “generations before present”. Users should keep in mind that ancient DNA variant calls tend to have a higher error rate, which has the potential to corrupt size history inferences. For example, when I first ran phlash on the Neanderthal and Denisovan genomes in the Wohns et al. data, it estimated a sharp, counterintuitive increase in *N*_*e*_ across all ancient populations roughly 100 generations before sample lifetime (Figure D.5). I then performed some exploratory analysis of the Wohns et al. call set, and found a roughly four-fold enrichment for C → T (or complementary G → A) mutations compared to other mutation types in the ancient samples (Figure 9a). This motif is known to arise in ancient DNA through a chemical process known as deamination (Dabney, Meyer, and Pääbo, 2013). Hence, the introduction of spurious variants can cause phlash to inflate estimates of *N*_*e*_ in the recent past.

**Figure 9:**
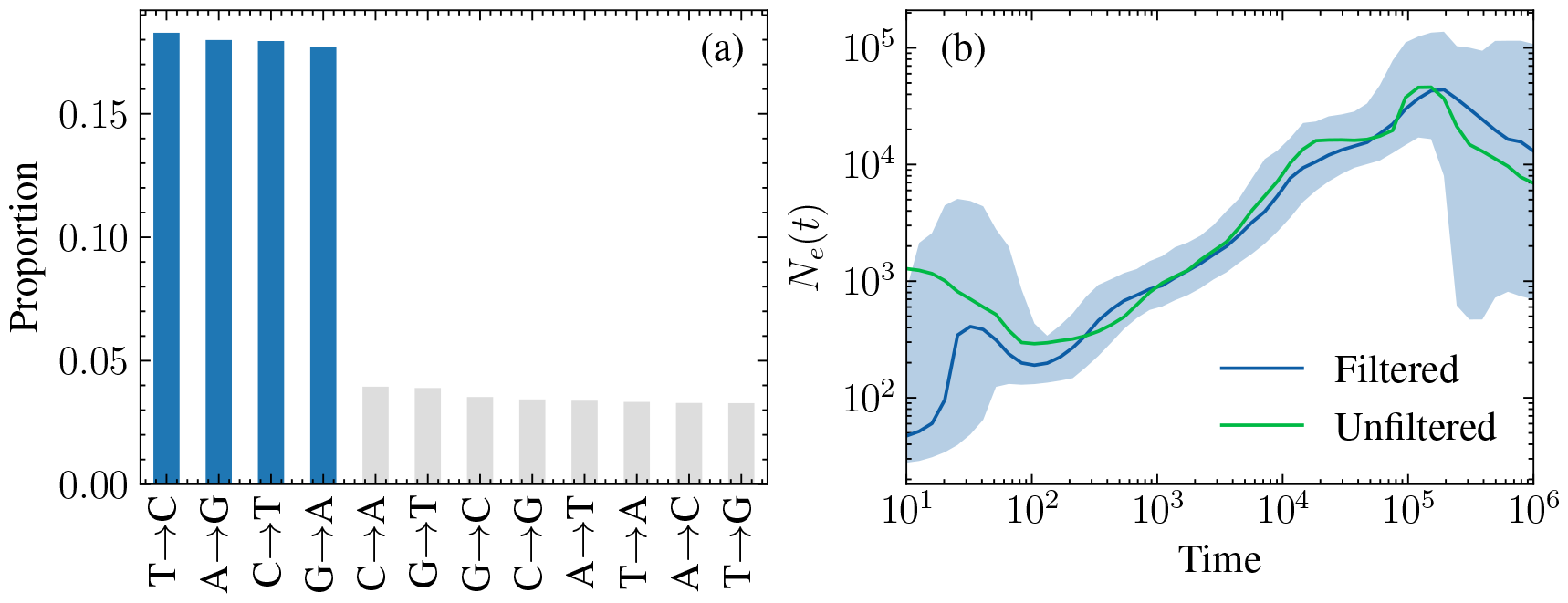
Analyzing ancient DNA using phlash. a) Histogram of the different mutation types in the ancient samples. Enrichment for C → T mutations are likely the result of deamination, as are T → C mutations after accounting for uncertainty in the ancestral allele. G ↔ A mutations are likely the result of deaminated bases on the complementary strand. b) Size history estimates for the Vindija Neanderthal before and after filtering out probable deaminated bases.

To correct for this effect, I re-ran phlash on a filtered version of the Wohns et al. call set which excluded all C ↔ T and G ↔ A variants that were private to the ancient populations. Filtered and unfiltered results are shown for the Vindija population in (Figure 9b) and for all ancient populations in Figure D.5. Filtering mostly removes the sudden, recent increase in *N*_*e*_, although evidence of it still persists to an extent in the credible intervals. It is possible that this simplistic filter does not detect other types of errors that can arise in aDNA. Users who want to study ancient samples using phlash, or other demographic inference tools, will want to pay careful attention to issues of data quality and filtering (see Orlando et al., 2021 for a review of best practices).

### 4.4 Detecting a population bottleneck

Finally, I studied the phlash’s ability to infer a sharp bottleneck in human data. Hu et al. (2023) have recently claimed that the population ancestral to all modern humans experienced an extreme bottleneck during the period 930–813kya, with the effective population size reduced to roughly 1,280 breeding individuals, or ∼1.3% of the ancestral size. The findings are based on fitting a novel demographic inference procedure, FitCoal, to frequency spectrum data from various African and non-African subpopulations in the 1000 Genomes and HGDP-CEPH panels. Curiously, although their method detected a bottleneck in African populations, it failed to do so in any of the 40 non-African populations surveyed, despite all extant populations having descended from a common basal population during the period in question (Bergström, Stringer, et al., 2021). Hu et al. attribute this discrepancy to the fact the out-of-Africa dispersal hinders the chance of discovering an ancient severe bottleneck, and suggest that in suitably corrected data, it is detectible in all modern populations. Finally, they assert that PSMC, SMC++, and RELATE (Speidel et al., 2019) all underestimated the severity of the ancient bottleneck.

I investigated whether phlash, which makes fewer parametric modeling assumptions and provides uncertainty quantification, could shed additional light on possibility of a human near-extinction event. First, I separately fit size history estimates to all of the distinct subpopulations in the Wohns et al. dataset, for a total of 159 different estimates. Posterior medians for each subpopulation are plotted in Figure 2(a), with the putative bottleneck interval shaded in gray (assuming a generation time of *g* = 24 years, as in Hu et al.). *A posteriori*, the data show little sign of a bottleneck, with all populations tightly clustered around *N*_*e*_ ≈1*3, 000* during the suggested interval. However, these estimates merely reflect the central tendency of each posterior—could stronger evidence emerge by examining the full distribution? I next considered distribution of the functional

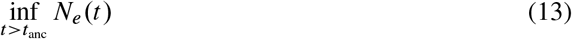

evaluated over all populations. Here, *t*_anc_ is a cutoff time designed to focus the estimator on ancient times (i.e., before the OOA bottleneck). I conservatively chose *t*_anc_ = (5 × 10^5^)/ *=* 24 generations, i.e. 500kya, which should give the estimator good power to detect a bottleneck in the distant past. A histogram of the posterior median of this functional is plotted in Figure 10(a). Although there is somewhat greater dispersion, there is still little evidence that *N*_*e*_ dipped below 10^4^ during the period in question.

**Figure 10:**
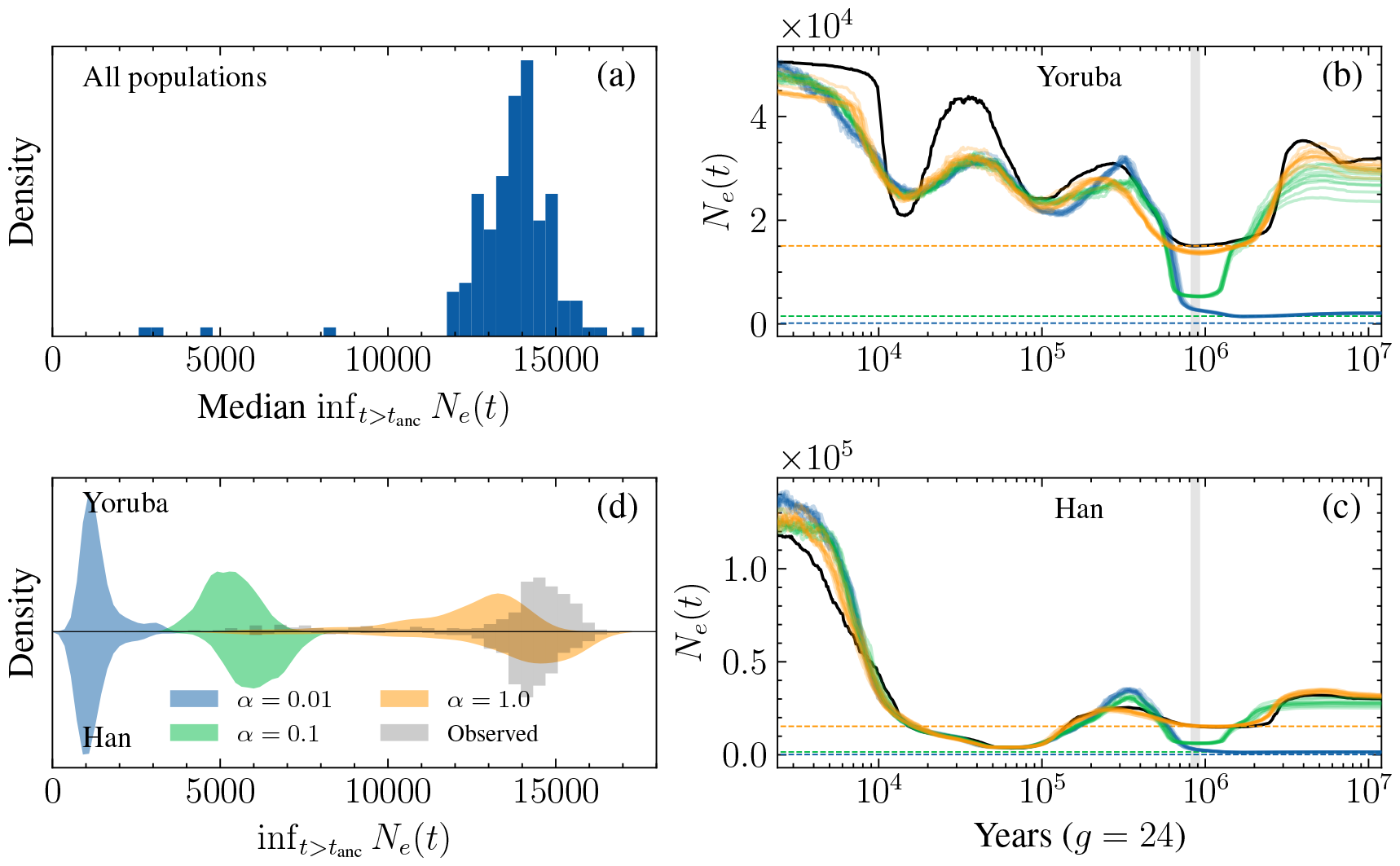
Searching for signs of an ancient bottleneck in real and simulated human data. (a) Histogram of posterior median of 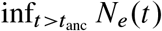 for each of the 159 populations considered. (b-c) The estimated size history of the Yoruba/Han population (black line, estimated from real data) was modified to incorporate a bottleneck. Orange, green, and blue correspond to bottleneck strength α ∈ {10^0^; 10^−1^; 10^−2^} respectively. Grey shaded region is [813 × 10^3^; 930 × 10^3^] years before present, assuming a generation time of 24 years. Dashed lines represent α*N*_*e*_(*t*), the population size during the bottleneck, and solid lines represent the corresponding phlash estimates. (d) Mirror plot of posterior density of test statistic for Han and Yoruba. Shaded density curves are its distribution in simulated data; bar plots are a histogram of its distribution in real data.

One potential explanation of these results is that phlash is biased, a possibility already suggested by Hu et al. with regard to the methods mentioned above. I view this as unlikely in light of the simulation results presented above; Figures D.1–D.4 show that phlash is quite generally quite accurate in the distant past, across a wide range of systems. Nevertheless, to probe further, I then followed Hu et al.’s methodology by simulating data under a pre-estimated model, introducing an artificial bottleneck, and then assessing accuracy. I performed this experiment for two populations, Han and Yoruba, that did and did not experience the OOA bottleneck, respectively. The number of diploid genomes for each population was *n*_Han_ = ∈48 and *n*_Yoruba_ = ∈∈4.

Using the fitted size histories for these two populations say, 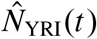 and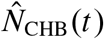, I introduced bottlenecks of strength α ∈ {1.*0, 0*.1, 0.01}, such that the perturbed size history was

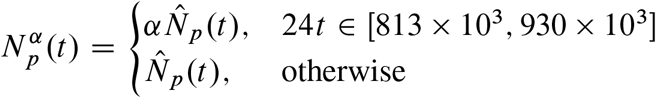

for *p* ∈ {YRI, CHB}. Results are shown in Figure 10(b)-(c): the black line is the baseline estimate, 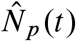, and the colored lines are the result of simulating from 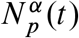 and then estimating to obtain 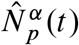. A total of ten replicates were performed for each setting of *p* and α, for a total of 60. The bottleneck interval is depicted in shaded gray. The α = 1 scenario (orange lines) reiterate that phlash has good power to estimate the true underlying size history function. When α = 0.1 (green lines), the phlash estimates dip noticeably, though some upward bias is apparent when comparing 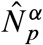 and 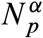 (the true level of 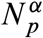 is shown as a dashed green line). Finally, when α = 0.01(blue lines), the phlash estimates closely track the true size history function, but fail to increase after the bottleneck interval, presumably because the severe bottleneck has forced all lineages to coalesce.

Focusing on the α = 0.01case, which corresponds to the result claimed by Hu et al., we can conclude the following: even though phlash does not perfectly recapitulate the size history for the period 24*t* > 930 × 10^3^, there is a substantial difference in the estimated size history functions between the α = 0.01 and α = 1 cases. In fact, as shown in Figure 10(d), the posterior distributions of the test statistic (13) are effectively disjoint between the two scenarios. From a decision-theoretic standpoint, this implies nearly perfect statistical power to distinguish the null hypothesis *H*_0_ : α = 1 from the alternative *H*_1_ : α = 0.01(e.g., Berger, 2013). In other words, if *H*_1_ were true, it would be readily discernible in data, irrespective of any estimation bias, because the test statistic would lay uniquely in the support of one of two disjoint distributions, and conversely for *H*_0_. In fact, this is precisely what I observed: the empirical distribution of the test statistic (Figure 10(d), grey bars) closely matches the α = 1 scenario for both populations, and barely overlaps with the α = 0.01 distribution at all. Thus, at least in the limited set of analyses I have performed here, I find little support for the strong ancient bottleneck hypothesis.

## 5 Conclusion

In this paper I have presented phlash, a new method for estimating historical effective population size. Using an extensive battery of simulation tests, I showed that phlash tends to be more accurate and efficient than several state-of-the-art methods. Moreover, the posterior distribution returned by phlash has a number of other uses, including uncertainty quantification, detection of population structure, and testing for ancient population bottlenecks. phlash is implemented as an efficient, open source Python package (see Data availability) and features a user-friendly API: almost all of the analyses presented here require but a few lines of code, and took under 60 minutes per population to run. One important caveat is that the implementation currently requires an Nvidia GPU. I hope to extend it it to work with AMD (and potentially other) accelerators in the future.

A few other possible extensions and avenues for improvement come to mind. Here I chose a simple prior model (equations 6–8) for *η*, which fixes the time discretization *t*_1_ < … < *t*_*M*_ to a logarithmically-spaced grid with random endpoints. A more flexible model would permit the time discretization to be completely arbitrary, which could allow for greater adaptivity. Indeed this was my initial approach, but I found that the particle-based sampling algorithm had difficulty converging, and so discarded it in favor of the simpler model with fewer parameters. Also, as noted in Section 3.1, phlash’s estimation error sometimes increased with larger sample sizes. I believe that this is because the composite likelihood (9) places equal weight on the “PSMC” and “SFS” components, which leads to the SFS component dominating for large *n*, combined with greater variance in the low-frequency SFS entries for large sample sizes. A smarter scheme might be to adaptively weight the the two terms, based perhaps on some measure of out-of-sample error, and to employ a binning procedure in the SFS component of the likelihood, as in Bhaskar, Y. X. R. Wang, and Y. S. Song (2015). Finally, phlash currently assumes a single global recombination rate parameter across all analyzed contigs. It could be generalized to learn contig-specific rates or, with considerably more effort, to utilize a fixed (non-learnable) position-specific rate map. Here I do note, however, that estimation quality seems to depend only very weakly on accurate knowledge of the recombination rate, as previously remarked by other authors (Schiffels and Durbin, 2014).

As far as extensions, the main technical contribution of the paper, which is a fast way to evaluate the gradient of the PSMC log-likelihood function, seems generally useful for population genetic inference. Indeed, the so called “inverse instantaneous coalescent rate” function, denoted *r*, in this paper, has been utilized in a number of previous studies to estimate more complex models than the simple panmictic one considered here (Rodríguez et al., 2018; Arredondo et al., 2021; Boitard et al., 2022; Mazet and Noûs, 2023). Extending these methods to take advantage of a differentiable likelihood function is, technically at least, straightforward. Similarly, certain methods which take as input pre-called identity-by-descent (IBD) tracts (e.g., Al-Asadi et al., 2019) can be re-cast as probabilistic models which depend on an underlying PSMC likelihood, and could be generalized to obtain procedures which integrate over all possible IBD scenarios, instead of fixing one of them *a priori*.

## Data availability

phlash is available as a Python package from https://github.com/jthlab/phlash. Code to reproduce the experiments is available at https://github.com/jthlab/phlash_paper. Tree sequences inferred by Wohns et al. (2022) containing the 1000 Genomes, HGDP, SGDP, and ancient genomes are available at https://zenodo.org/records/5512994.

## Acknowledgements

I am grateful to Dat Do for helpful providing comments on a draft of the manuscript. This research was supported by NSF grant DMS-2052653 and the National Institute of General Medical Sciences of the NIH under award number R35GM151145. The content is solely the responsibility of the author and does not necessarily represent the official views of the NIH.

## A Fast computation of the score function

This section describes a fast algorithm for computing the score function of the psmc model. The algorithm combines several existing results from the literature. The first of these is Fisher’s identity (Cappé and Moulines, 2005), which is a general expression for the score function of a latent variable model. To state the identity I recall some notation from their paper. Let 0 be a parameter, *q*^θ^ (*x, x*′) denote the transition function between states *x* and *x*′ in a hidden Markov model, and 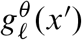 denote the probability of the 𝓁-th observation when the hidden state at position 𝓁 is *x*′. Define *r*_0_(*x, x*′) = *π*(*x*)*q*^θ^ (*x, x*′)*g*_1_(*x*′), where *π* is a prior on the initial state of the hidden Markov chain, and *r*𝓁 (*x, x*′) = *q*^θ^ (*x, x*′)*g*_𝓁+1_(*x*′) for *k* > 0. Fisher’s identity applied to the hidden Markov model is then

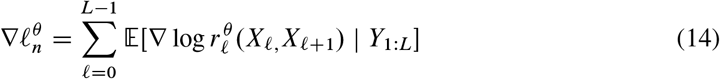

where the expectation is with respect to the latent states *X*_0:*L*_ given the observed data *Y*_1:*L*_. This identity gives the intuitive result that the score function decomposes as a sum of expectations over the pairwise posterior distributions (*X*_𝓁_, *X*_𝓁+1_) | *Y*_1:*L*_. It is well known that these posterior distributions may can recursively computed using the forward-backward algorithm; indeed this is essentially the “E”-step of the EM algorithm used by for parameter estimation in HMMs (Bishop, 2006). However, this requires storing the results of the forward pass which costs 𝒪(*LM*) memory which, as noted above, is prohibitive in the GPU setting.

In fact, there is a less well-known variant of the Baum-Welch algorithm which has lower memory and higher computational cost. Jensen (2009) shows how the so-called “linear memory” Baum-Welch algorithm can recursively compute posterior expectations of the form

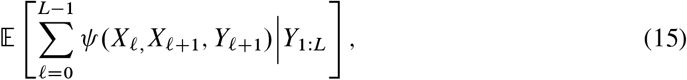

where *ψ* (*x, x*′, *y*) is an arbitrary functional, using only a single pass over the data. Observe that (14) has the form (15) for 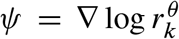. Therefore, the linear memory Baum-Welch algorithm can also be used to compute the score function. A single run of the algorithm takes 𝒪(*LM* ^2^) time. Hence, to run the standard Baum-Welch algorithm using the linear-memory variant, we would have to compute (15) for ψ (*x, x*′, *y*) = **1**{*x* = *i, x*′ = *j* } for all *i, j* ∈ [*M]*, at an overall cost of 𝒪(*LM*) operations, explaining why the algorithm is not widely used in practice.

### A.1 A constant memory quadratic-time algorithm

In this section I show how the linear memory Baum-Welch algorithm can be combined with fast matrix-vector multiplication algorithms that have been developed for coalescent hidden Markov models to yield a new algorithm that is simultaneously time- and memory-efficient. The central recursive quantity analyzed by Jensen (2009) is

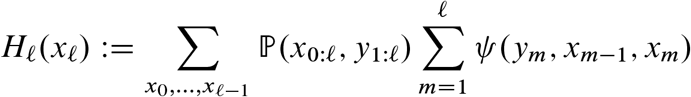

which is (proportional to) the posterior expectation of 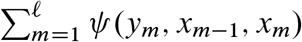 evaluated over all hidden state paths ending in *X*_𝓁_ = *x*_𝓁_. Jensen shows that *H*_𝓁_(*x*_𝓁_) satisfies the recursion

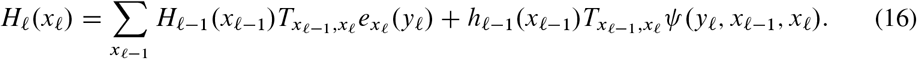

where 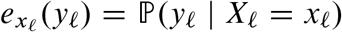 if the emission probability of the symbol observed at position 𝓁 conditional on the latent state being *x*_𝓁_, and *h*_𝓁_(*x*_𝓁_) = P (*x*_𝓁_, *y*_1:𝓁_) is the filtering density.

To advance the algorithm one step we have to compute *H*_𝓁_(*x*_𝓁_) for all *x*_𝓁_ ∈ {1, …, *M* }. Because of the summation in (16), computing each entry costs 𝒪(*M*) FLOPS, for a total cost of 𝒪(*M* ^2^). Note also that once *H*_𝓁_ is computed, the algorithm no longer makes use of *H*_*i*_ for *i* < 𝓁. Hence the memory requirement is only 𝒪(*M*) per computed φ; in particular, it does not depend on the sequence length *L*.

To arrive at a faster algorithm I first observe that (16) can be written more compactly in matrix notation as

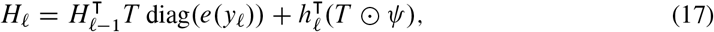

where ⊙ denotes Hadamard (entry-wise) product. Now, the term 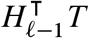 is a standard matrix-vector multiply costing 𝒪(*M* ^2^). However, in the specific case where *T* is the transition matrix of a psmc-like hidden Markov model, Palamara et al. (2018) show that matrix-vector products of the form *v*^Τ^*T* can be computed in 𝒪(*M*) time. (Note that, although *v* is a probability vector in their application, the result holds for an arbitrary *v* ∈ ℝ^*M*^ .) Hence, the cost of computing the first term in (17) can immediately be reduced to 𝒪(*M*).

The second term, 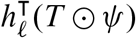, does not necessarily have the structure needed for the linear-time matrix-vector multiply algorithm to work. However, by reparameterizing the transition matrix of the SMC coalescent hidden HMM in a favorable way, I can ensure that it can be computed in time linear in *M*. The next section describes how to do that.

### A.2 Structure of the SMC’ transition matrix

Palamara et al. (2018) have shown that the transition matrix for the SMC, SMC’, and conditional Simonsen-Churchill (CSC) models have the following structure:

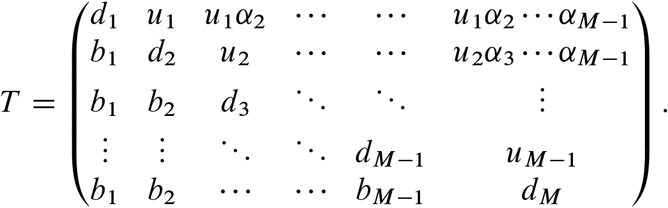

(Here *T* is a discretized version of the corresponding transition function *q*(*x*_𝓁_ | *x*_𝓁-1_, *r*,) defined in equation (2).) In the above display, *b, u* ∈ ℝ^*M*-1^ are the first sub- and super-diagonals, respectively, *d* ∈ ℝ^*M*^ is the main diagonal, and α_*j*_ = *T*_1,*j*+1_*=T*_1,*j*_ is the ratio of successive columns in the first row.

I first reparametrize this matrix in a different coordinate system which will make computation of score function easier:

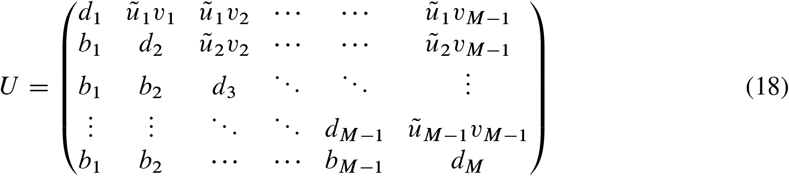

where I defined 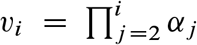 and 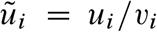. Now let *U* = log *T* denote the entry-wise logarithm of *T* in the above parameterization. We then have

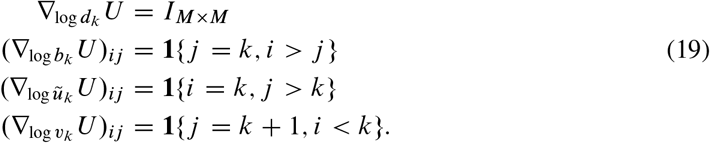

Now let 𝓁 denote the log-likelihood under the PSMC, PSMC’, or PSMC-CSC models. Consider how to compute ∂𝓁*=*∂ log *d*_*k*_ where *d*_*k*_ are the diagonal entries of the transition matrix *U*. Applying Fisher’s identity (14), we have

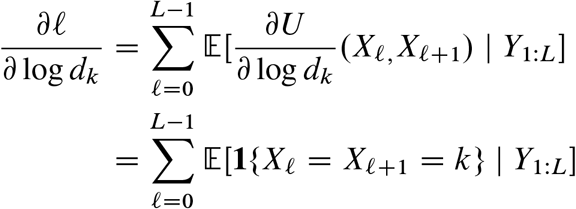

which is (15) with φ(*x*_𝓁_, *x*_𝓁_ + 1, *y*_𝓁+1_) = **1**{*x*_𝓁_ = *x*_𝓁_ + 1 = *k*}. Hence, one can apply the algorithm of the preceding section, where I note that, in this case,

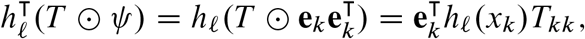

where **e**_*i*_ ∈ ℝ^*M*^ denote the standard coordinate vectors. Note that this computation takes 𝒪(*M*) time. Overall, computing 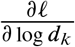 therefore requires 𝒪(*LM*) time and 𝒪(*M*) memory, and hence 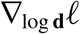 can be evaluated in 𝒪(*LM* ^2^) time and 𝒪(*M* ^2^) memory.

The score function with respect to the other entries of *U* can be computed in a similarly, if slightly more involved, manner. I show how to compute 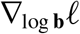 and omit the similar derivations of the remaining parameters. Using (19), we have

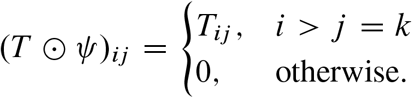

So *T* ⨀ ψ contains the lower diagonal entries of the *k*-th column of *T*, and is zero everywhere else. But by (18), the lower diagonal entries are constant, so that

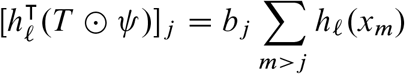

which can again be computed in 𝒪(*M*) time using recursion.

### A.3 Technical considerations

One important caveat of the algorithm is that, for technical reasons related to the Nvidia CUDA architecture, performance is currently optimized when *M* = 16, or a small multiple thereof. This occurs for two reasons: a) the size of a half-warp on all existing CUDA GPUs is 16 threads, and b) the memory requirement of the algorithm is roughly 28*M* ^2^ bytes. With *M* = 16, the state of the entire algorithm can be store in on-chip shared memory, which is hundreds of times faster to access that global/hierarchical block memory.

Although *M* = 16 results in a somewhat coarse approximation, the experimental results in the main text show that this choice does not overly bias the estimates. Furthermore, it should be possible to increase *M* with future generations of hardware.

## B Algorithm for computing credible bands

Given a posterior sample 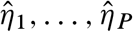 from ℙ (*η* | **G**), let

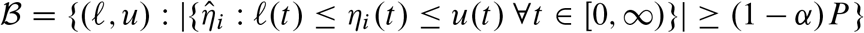

denote the set of all bracketing functions (𝓁, *u*) that contain at least a 1 - α fraction of the sample. The (1 - α) credible band is the function pair (𝓁^*^, *u*^*^) such that, for any (𝓁, *u*) ∈, there exists a *t* ∈ [0, ∞) such that 𝓁(*t*) ≤ 𝓁^*^(*t*) or *u*(*t*)≥ *u*^*^(*t*); in other words, the tightest possible band in. The credible band can be determined by solving a mixed-integer linear programming problem.

Let 0 = *t*_0_ < *t*_1_ < · · · < *t*_*K*_ be a grid of timepoints such that 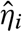 is constant on [*t*_*j*_, *t*_*j*+1_) for all *i* and *j* ; this is always possible for large enough *K* since each 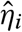 is piecewise-constant. Then define the decision variables

- *u*_*k*_: upper bound on the confidence band at *t*_*k*_.
- *𝓁*_*k*_: lower bound on the confidence band at *t*_*k*_.
- *z*_*ik*_: binary indicator that equals 1 iff 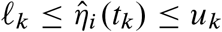.

The objective function is

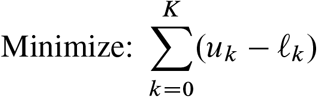

subject to the constraints:

- 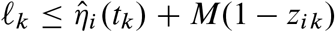, for all *i* and *k*,
- 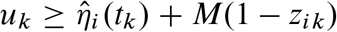, for all *i* and *k*,
- 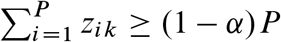, for all *k*,
- *𝓁*_*k*_, *u*_*k*_ ≥ 0, for all *k*.

where *M* is a sufficiently large constant such that the constraint is ignored if *z*_*ik*_ = 0 (the so-called “big *M*” method).

Exactly solving the optimization problem amounts to taking the union of all time points in all of the models *η*_*i*_, which are almost surely distinct, so that *K* = 16 × 500 using the program defaults. This results in a very large optimization problem which takes many days to solve. For this reason, the implementation in phlash defaults to approximating each 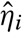 at a sparse grid of points. Specifically, it finds 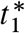 and 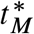 such that the posterior sample is constant outside of 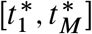 and then approximates each function at a geometrically spaced grid of 100 points 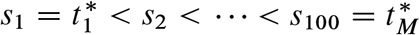,

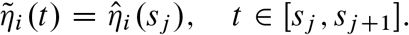

The optimization problem is then solved with 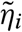 in place of 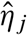.

## C Command lines

The command line used to run `scrm` was:

~~~
scrm {sample size} 1 {demographic model} -t {theta} -r {rho} {L}
 -transpose-segsites -SC abs -p 14 -oSFS -seed {seed}
~~~

where `<sample size>`, `theta`, etc. denote various simulation- and species-specific parameters. The demographic model parameters were obtained by exporting each `stdpopsim` model into a sequence of `ms` demography commands using `demes` (Gower et al., 2022).

The command line use to run SMC++ was:

~~~
smc++ estimate --cores 4 --knots 24 -o {outdir} {mutation_rate} {input_files}
~~~

The command line used to run MSMC2 was:

~~~
msmc2_Linux -o {outdir}/output -I {pairs} -t4 {hets}
~~~

where `pairs` was set equal to the sequence {(2*i*,2*i* + 1) : *i* ∈ {1, …, *n*}}. This setting ensured that MSMC2 only had access to unphased data, and also reduced memory consumption.

The command line used to run FITCOAL was

~~~
java -cp {jar} FitCoal.calculate.SinglePopDecoder
-table {tables}
-input {afs} -output {output_base}
-generationTime 1
-mutationRate {mutation_rate_per_kb}
-omitEndSFS {trunc}
-randSeed {seed}
-genomeLength {genome_length_kbp}
~~~

The `trunc` parameter in the preceding display was obtained by first running

~~~
java -cp FitCoal.jar Fitcoal.TruncateSFS
~~~

as suggested in the FITCOAL README.

## Additional Figures and Tables

**Table D.1:** Description of the simulation models used to benchmark each method. Each model is taken from the `stdpopsim` catalog (Adrion et al., 2020). *r* is the recombination rate, computed as a weighted average over the chromosome-specific rates. *L* is total genome length in gigabase-pairs; simulations were restricted to recombining autosomes.

| | Species | Model | Population | Description | $\mu$/bp/gen | $r$/bp/gen | $L$ (Gbp) |
| --- | --- | --- | --- | --- | --- | --- | --- |
| 1 | Anopheles gambiae | GabonAg1000G_1A17 | GAS | Stairwayplot estimates of $N(t)$ for Gabon sample | $3.5 \times 10^{-9}$ | $1.38 \times 10^{-8}$ | 0.206 |
| 2 | A. thaliana | African3Epoch_1H18 | SouthMiddleAtlas | South Middle Atlas African three epoch model | $7 \times 10^{-9}$ | $8.06 \times 10^{-10}$ | 0.119 |
| 3 | A. thaliana | SouthMiddleAtlas_1D17 | SouthMiddleAtlas | South Middle Atlas piecewise constant size | $7 \times 10^{-9}$ | $8.06 \times 10^{-10}$ | 0.119 |
| 4 | Cattle | HolsteinFriesian_1M13 | Holstein_Friesian | Piecewise size changes in Holstein-Friesian cattle. | $1.2 \times 10^{-8}$ | $9.26 \times 10^{-9}$ | 2.49 |
| 5 | D. melanogaster | African3Epoch_1S16 | AFR | Three epoch African population | $5.49 \times 10^{-9}$ | $2.01 \times 10^{-8}$ | 0.109 |
| 6 | Human | Africa_1T12 | AFR | African population | $1.29 \times 10^{-8}$ | $1.28 \times 10^{-8}$ | 2.88 |
| 7 | Human | AmericanAdmixture_4B11 | ADMIX | American admixture | $1.29 \times 10^{-8}$ | $1.28 \times 10^{-8}$ | 2.88 |
| 8 | Human | Constant | pop_0 | Constant population size, $N_e = 10^4$. | $1.29 \times 10^{-8}$ | $1.28 \times 10^{-8}$ | 2.88 |
| 9 | Human | Zigzag_1S14 | generic | Periodic growth and decline. | $1.29 \times 10^{-8}$ | $1.28 \times 10^{-8}$ | 2.88 |
| 10 | Chimpanzee | BonoboGhost_4K19 | bonobo | Ghost admixture into bonobos | $1.6 \times 10^{-8}$ | $1.2 \times 10^{-8}$ | 2.79 |
| 11 | Olive baboon | SinglePopSMCpp_1W22 | PAnubis_SNPRC | SMC++ estimates of $N(t)$ for Papio Anubis individuals | $5.7 \times 10^{-9}$ | $1.12 \times 10^{-8}$ | 2.59 |
| 12 | Sumatran orangutan | TwoSpecies_2L11 | Bornean | Two population orangutan model | $1.5 \times 10^{-8}$ | $6.4 \times 10^{-9}$ | 2.67 |

**Figure D.1:**
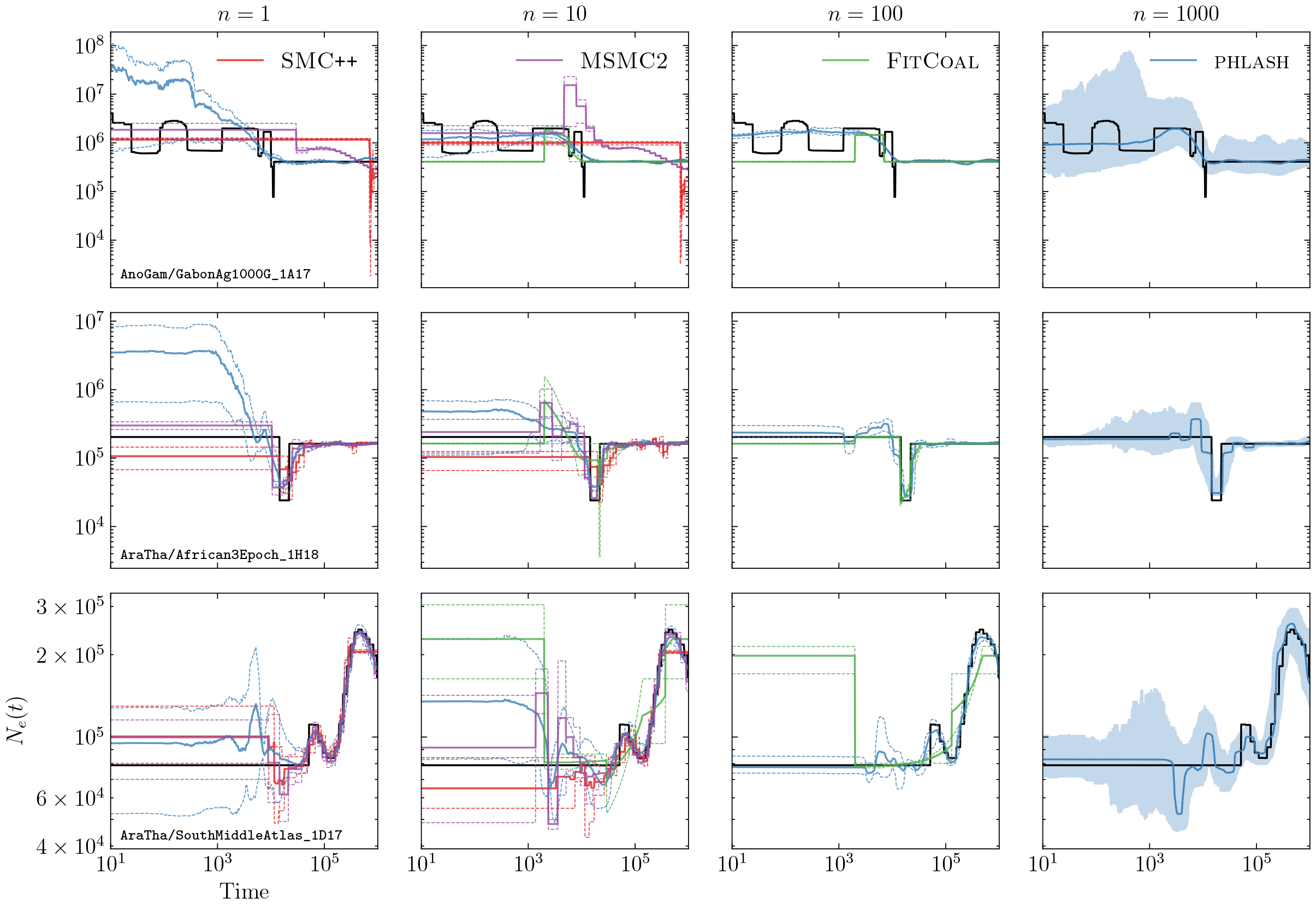
Simulation results for models 1–3. For *n* = 1000, shaded regions indicate 95% credible intervals. For all other *n*, the solid (dotted) lines represent the median (min/max) across all simulation replicates.

**Figure D.2:**
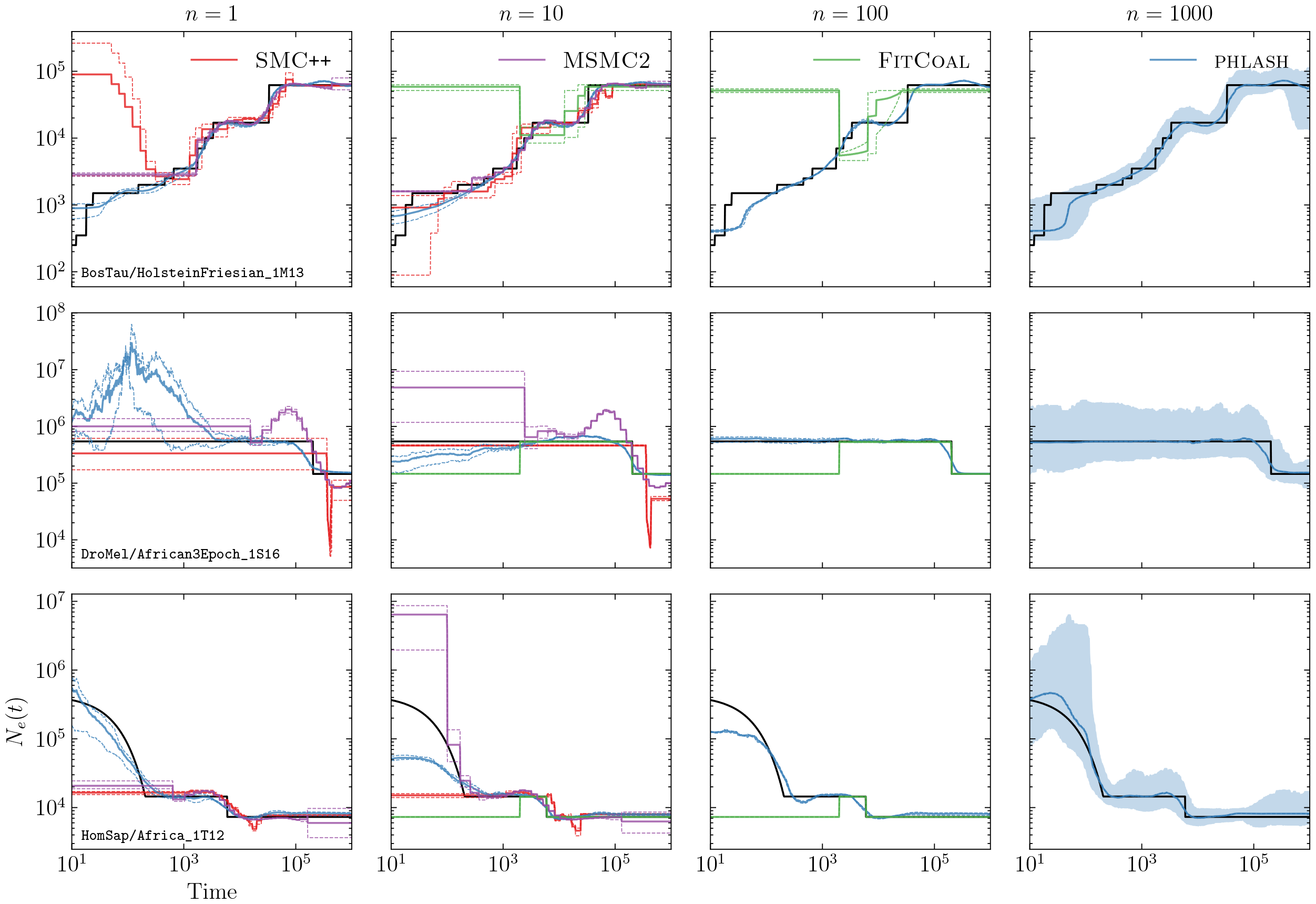
Simulation results for models 4–6. See Figure D.1 for additional information.

**Figure D.3:**
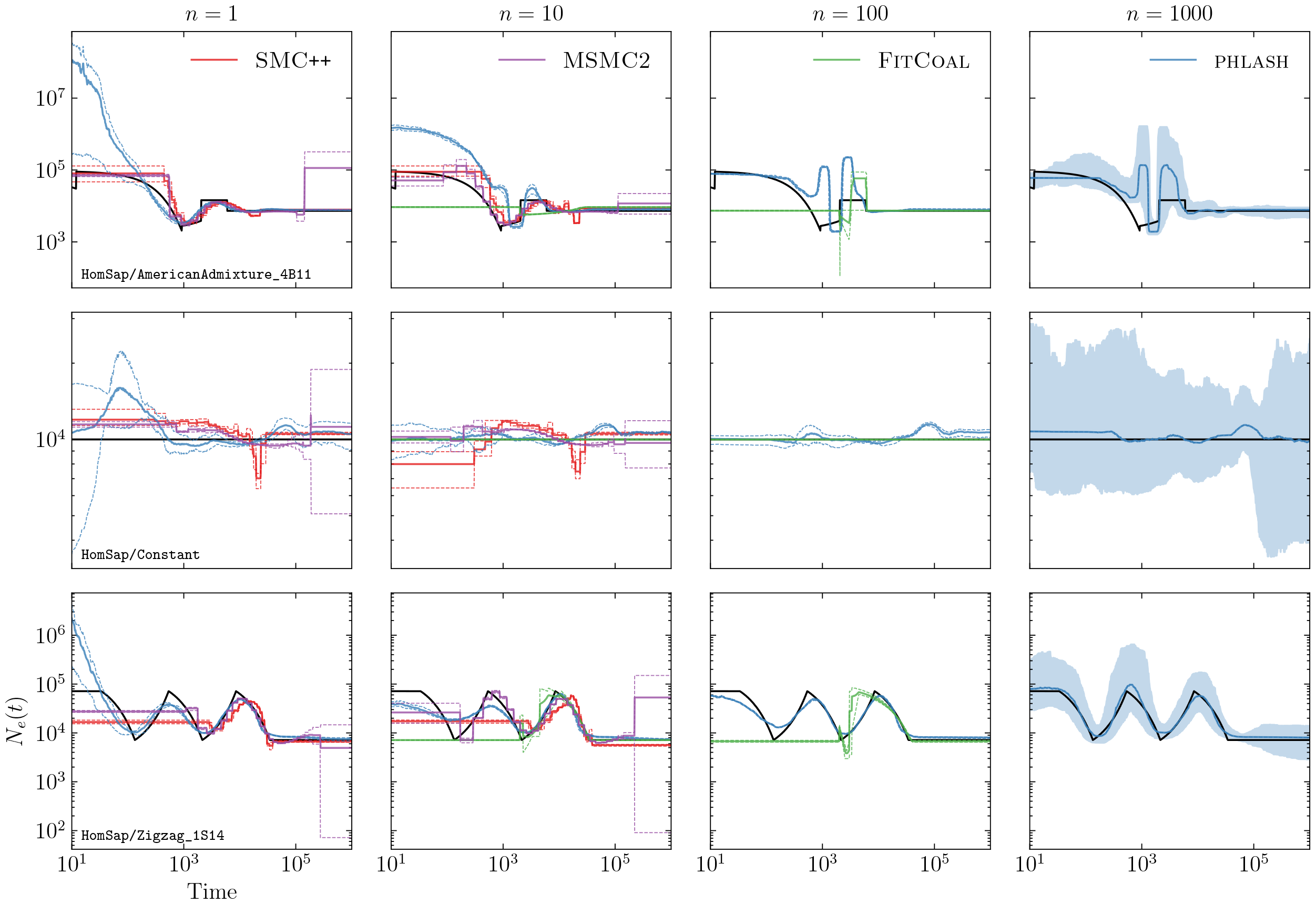
Simulation results for models 7–9. See Figure D.1 for additional information.

**Figure D.4:**
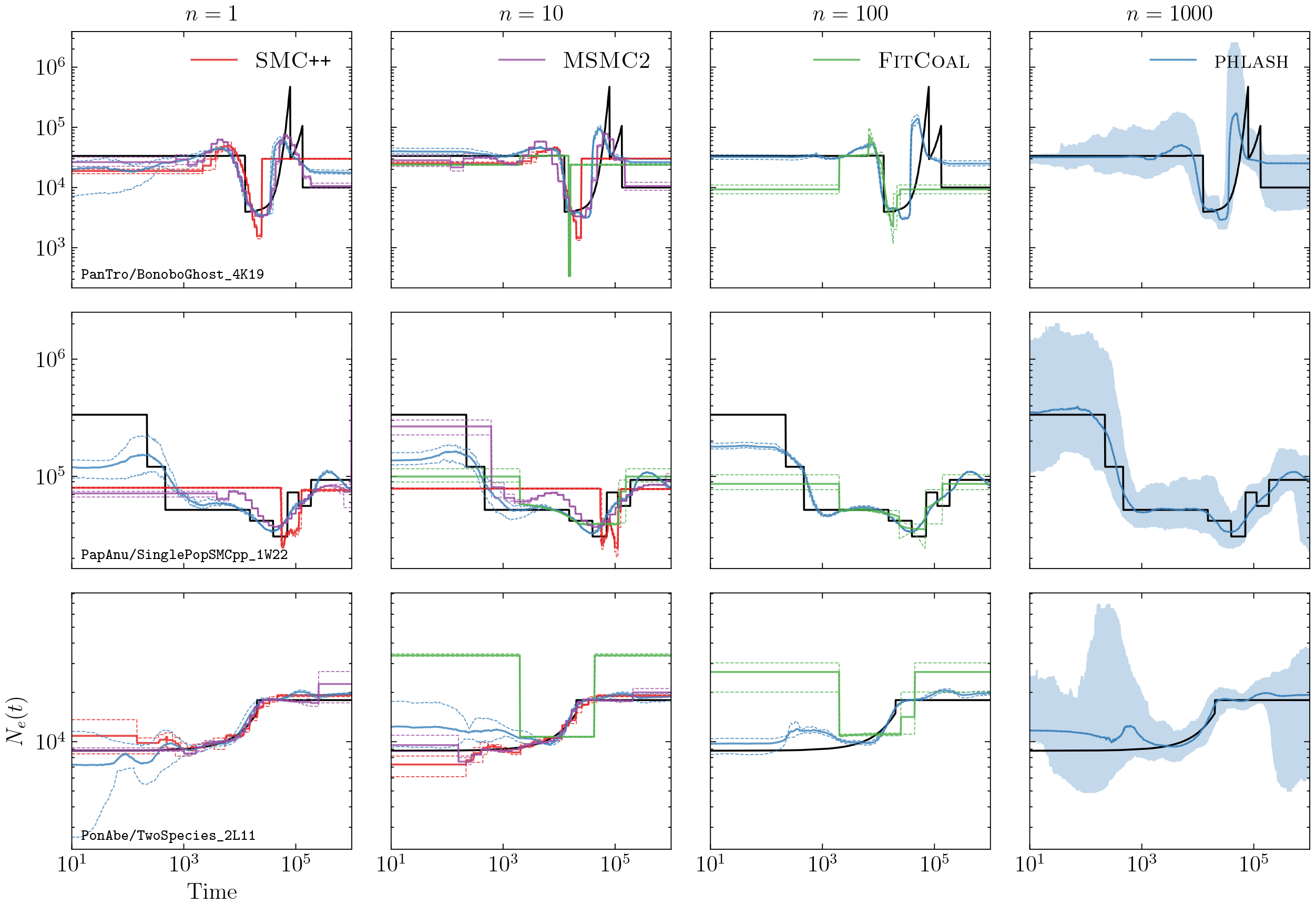
Simulation results for models 10–12. See Figure D.1 for additional information.

**Figure D.5:**
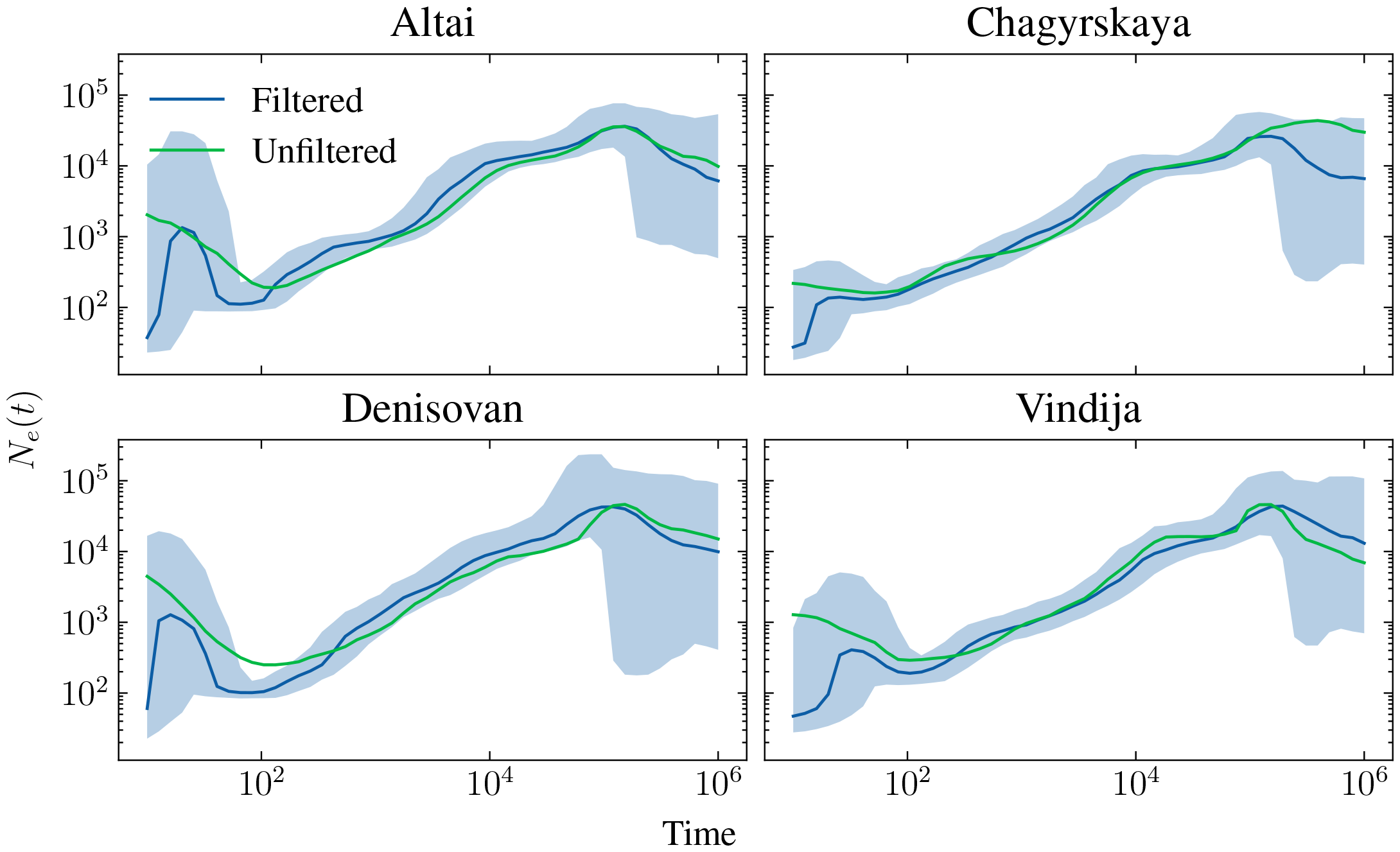
*N*_*e*_ estimates for archaic humans. Solid lines are posterior medians, and blue shaded area shows the 95% credible intervals for the filtered data. “Unfiltered” refers to the raw genotypes extracted from the Wohns et al. (2022) dataset, while “filtered” are the estimates obtained after running the filtering procedure described in the main text.

